# Dynamics of task-related electrophysiological networks: a benchmarking study

**DOI:** 10.1101/2020.08.02.232702

**Authors:** Judie Tabbal, Aya Kabbara, Mohamad Khalil, Pascal Benquet, Mahmoud Hassan

## Abstract

Motor, sensory and cognitive functions rely on dynamic reshaping of functional brain networks. Tracking these rapid changes is crucial to understand information processing in the brain, but challenging due to the random selection of methods and the limited evaluation studies. Using Magnetoencephalography (MEG) combined with Source Separation (SS) methods, we present an integrated framework to track fast dynamics of electrophysiological brain networks. We evaluate nine SS methods applied to three independent MEG databases (N=95) during motor and memory tasks. We report differences between these methods at the group and subject level. We show that the independent component analysis (ICA)-based methods and especially those exploring high order statistics are the most efficient, in terms of spatiotemporal accuracy and subject-level analysis. We seek to help researchers in choosing objectively the appropriate methodology when tracking fast reconfiguration of functional brain networks, due to its enormous benefits in cognitive and clinical neuroscience.

## Introduction

Evolving evidence show that motor, sensory, emotional and cognitive functions emerge from dynamic interactions between cortical and subcortical brain structures. Specific rhythms of neural networks allow synchronization and long-range communication between distant and distributed brain areas. This phenomena was shown crucial during visual (Bola and Sabel, 2015; Hassan et al., 2015; Mheich et al., 2018), auditory (Fontolan et al., 2014), sensorimotor (Pomper et al., 2015; Wilkins and Yao, 2020) and cognitive (Negrón-Oyarzo et al., 2018; Rouhinen et al., 2020) tasks. This brain communication is very transient and there is a dynamic reorganization of functional brain networks during behavioral tasks, even at sub-second time scale (Vidaurre et al., 2018b). Therefore, the analysis of whole-brain dynamic functional connectivity (dFC) has become a burgeoning field of research in cognitive neuroscience (Bassett and Sporns, 2017; Bullmore and Sporns, 2009; Iraji et al., 2020; Kabbara et al., 2020). In this regard, Magneto/Electro-encephalography (MEG/EEG) provides a unique direct and noninvasive access to the electrophysiological activity of the whole brain, at the millisecond scale. Benefiting from the excellent time resolution of the MEG/EEG (~millisecond), current methods allow of estimating sub-second time-varying functional brain networks in the cortical space through sensor-level signals (Hassan et al., 2014; Hassan and Wendling, 2018). The key challenge here is how to characterize and quantify these rapidly changing networks.

Several frameworks have been used and one can distinguish between approaches that analyze the time-varying signal such as data-driven techniques and those that more explicitly model/capture dynamics over time such as Hidden Markov Model (HMM) (Baker et al., 2014; Vidaurre et al., 2018a, 2018b, 2016), Autoregressive model (AR) (Casorso et al., 2019) and General Linear Model (GLM) (Friston, 1994). The data-driven methods, where ‘brain network states’ are derived directly from the data without a priori hypothesis on the fitting model, have showed promising results, despite the fact that the selection of the used algorithm is largely empirical. In these methods, sliding window approach, that forms a series of temporal networks, is usually used and followed by a dimensionality reduction or clustering approach. This includes, Kmeans(Allen et al., 2014; Ciric et al., 2017; Du et al., 2016; Fong et al., 2019; Liu and Duyn, 2013; Mheich et al., 2015; O’Neill et al., 2015), component analysis such as temporal Independent Component Analysis (tICA)(O’Neill et al., 2017), Principal Component Analysis (PCA)(Leonardi, n.d.) and Nonnegative Matrix Factorization (NMF)(Chai et al., 2017). Although the conceptual difference between these methods (and within each family of methods such as different ICA algorithms) is theoretically obvious (as they are based on different assumptions), the studies that investigate the differences between them remained very few. The existing comparative studies are mainly limited to confirming results of differences between two conditions(Leonardi, n.d.) or to prove that obtained results are unaffected by the method’s choice(Miller et al., 2016). However, a throughout quantitative and qualitative comparative study using both simulation-based and data-driven approaches is still missing and there is no clear consensus about the ‘best’ (if any) source separation or clustering method to be used to adequately tracking dFC, which is the main objective of our study.

Here, we evaluate the performance of nine dimensionality reduction methods used to track functional connectivity states at both group and individual levels. This was done using simulations and three independent MEG datasets (N=95) recorded during motor and working memory tasks (see Figure 1). The dynamic brain networks were reconstructed using MEG source connectivity method combined with a sliding window technique. The dimensionality reduction algorithms were compared in terms of their temporal and spatial accuracy. These methods include PCA, NMF, Kmeans and six various versions of ICA (Joint Approximation Diagonalization of Eigen-Matrices (JADE), INFOMAX, Second-Order Blind Identification (SOBI), fixed-point algorithm (FastICA), COM2 and Penalized Semi-Algebraic Unitary Deflation (P-SAUD). The motivation behind using several ICA subtypes is that each one has its own definition of statistical independence and several studies showed conceptual differences between them (Kachenoura et al., 2008; Sahonero-Alvarez and Calderon, 2017). We also analyzed the optimal number of subjects needed for each method to reveal significant results. Overall, our results show that the ICA methods and essentially those based on high order statistics have the highest spatiotemporal accuracy at group and subject level. This study aims at providing a well-established framework for researchers interested in studying reconfiguration of functional brain networks during cognitive processes.

**Figure 1.**
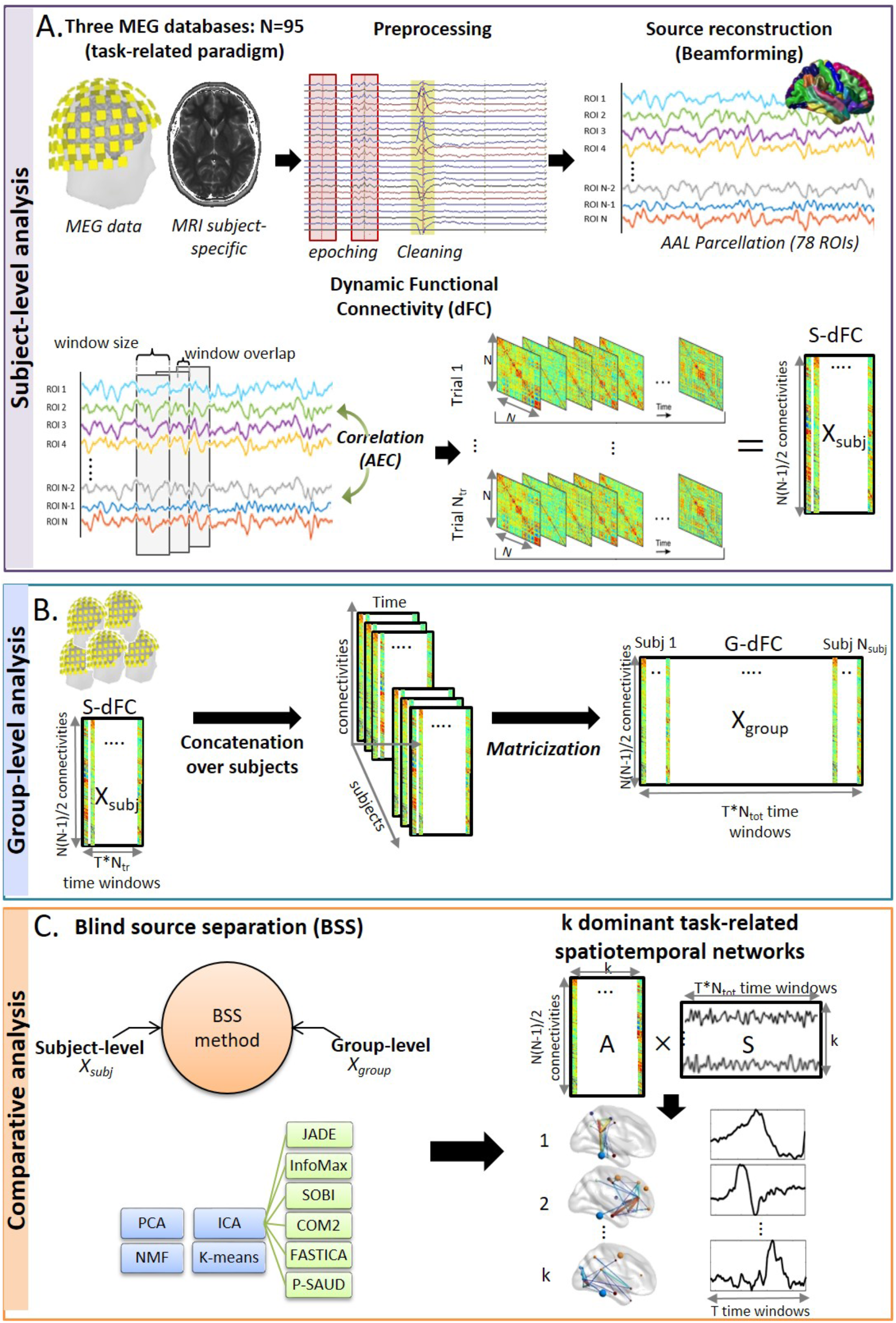
Illustration of the investigation structure for each of the three task-related paradigms. A. The fundamental processing pipeline applied on each subject data from sensor-level (using non-invasive MEG technique) to cortical-level (using beamforming as the inverse problem solution) to dynamic functional connectivity computation (S-dFC) (using the sliding window approach). B. Concatenation of S-dFC of all subjects along time axis to form a group data referred to as G-dFC, C. Comparative analysis between nine different source separation (SS) methods (six variants of ICA, PCA, NMF and Kmeans) applied on both group-level (X_group_) and subject-level (X_subj_) data in order to derive k dominant task-related spatiotemporal components (the mixing matrices represent brain spatial maps while the extracted sources represent corresponding temporal weights fluctuations).

## Materials and methods

### 1. Data

#### 1.1. Dataset1: ‘Self-paced button press task’

This dataset includes 15 healthy righthanded participants (9 male and 6 female, aged 25 ± 4 years (mean ± SD)). They were asked to press a button with the index finger of their non-dominant hand, once every 30 seconds, and should not count the time between presses. More details about this dataset can be found in (Kabbara et al., 2019; O’Neill et al., 2017).

#### 1.2. Dataset2: ‘HCP left hand movement Task’

61 healthy participants (28male and 33 female, aged 22-35) completed the MEG Motor task provided by the Human Connectome Project (HCP) (MEG-1 release)(Van Essen et al., 2012). The corresponding experimental protocol was adapted from Buckner and colleagues(Buckner et al., 2011; Thomas Yeo et al., 2011). It was performed in two sessions of 14min each, with a small break between them. Each session consisted of 42 total blocks randomly distributed; 32 of them were partitioned into 16 hand movements blocks (8 right and 8 left), and 16-foot movements blocks (8 right and 8 left), and the remaining 10 blocks were interleaved resting/fixation blocks. Each motor effector block was preceded by a 3sec visual cue that prompts participants to either tap their left or right index and thumb fingers or squeeze their left or right toes. The block lasted for 12sec and consisted of 10 sequential movements, each initiated with 150ms pacing stimuli followed by 1050ms black screen for task execution. Here, for simplicity, we were interested in the trials related to the left hand moves only. MEG data was recorded at Saint Louis University at 508.6275Hz sampling frequency and co-registered with the available subject specific MRI. EMG activity was also recorded from each limb.

#### 1.3. Dataset3: ‘Sternberg Working Memory Task’

19 healthy participants (10 male and 9 female, aged 25±3 years (mean ± SD)) performed Sternberg task, in which two example visual stimuli, mainly abstract geometric shapes, were successively presented on a screen; each for 0.6sec and separated by 1sec. Then, a maintenance period of 7sec was left before the presentation of a third probe stimulus. Consequently, subjects were asked to press a button with their right index finger only if the probe stimulus matched either of the two example stimuli and an immediate feedback will be given to show their response correctness. 30 trials were presented separated by 30sec of rest. In both datasets 1 and 3, MEG data were recorded using a 275-channel CTF MEG system at 600Hz sampling frequency and co-registered with subject-specific MRI. Both datasets were approved by the University of Nottingham Medical School Research Ethics Committee(O’Neill et al., 2017; Vidaurre et al., 2018a).

### 2. Methodology

#### 2.1. Preprocessing

Both datasets 1 and 3 were received already preprocessed as described in(O’Neill et al., 2017). Briefly, bad segments produced by muscles, eye or head movement were already visually inspected and removed. For dataset 2, we used the preprocessing pipeline offered by the HCP consortium, which includes removing bad channels, segments and bad independent components from task data. Segments were retrieved from the dataset 1 in the interval [-15; +15sec] relative to the button press onset, and from the dataset 3 in the interval of [-16; +28sec] relative to stimulus presentation. In HCP analysis, we chose data epochs time-locked to EMG onset as we were concerned in exploring brain networks involved during movement execution. Thus, trials were segmented in [-1.2; +1.2sec] relative to EMG onset. Then, as functional connectivity was proved to be frequency-dependent(Baker et al., 2014; Hipp et al., 2012), each dataset was preprocessed in its appropriate frequency band actively involved in the corresponding cognitive task. While beta band [13-30Hz] was used for self-paced motor task, working memory data was filtered in a broader band [4-30Hz] as it is has been shown to involve multiple frequency bands, according to previous studies(Brookes et al., 2012; O’Neill et al., 2017). Higher gamma frequencies [30-100Hz] were used for HCP data that deals with very rapid time scale tasks. After these preprocessing steps, an average of 34, 150 and 29 per subject were kept from dataset 1, 2 and 3, respectively.

#### 2.2. Source Reconstruction and Functional Connectivity

In order to localize brain sources and reconstruct their activities, we used the Linearly Constrained Minimum Variance Beamforming (LCMV) approach on parcellated cortex using AAL atlas (N=78 regions of interests -ROIs-). Following this, we estimated the functional connectivity by computing the amplitude envelope correlations (using Hilbert transformation) between all ROIs. Symmetric orthogonalization was also performed to remove signal leakage for both datasets 1 and 3 as recommended by (Colclough et al., 2015).

#### 2.3. Dynamic Functional Connectivity Analysis (dFC)

To estimate the dynamic functional brain networks, we adopt the widely used approach of sliding windows for all datasets. To this end, a time window of length 6sec with 0.5sec was used for datasets 1 and 3 as applied by Oneill et al.(O’Neill et al., 2017). Concerning the dataset 3, we used a 0.6sec length window with 0.05sec step to have similar range of number of windows obtained for all datasets. As a result, we obtained, for each subject trial, a ‘dynamic functional connectivity (dFC)’ matrix of dimension [NxNxT], where T=49, 36 and 75 for datasets 1, 2 and 3 respectively. Next, due to symmetry, we unfolded this matrix into a 2-D [Nx(N-1)/2×T] matrix by removing the redundant connections in each time window. Then, the mean of each row of this matrix is subtracted from the data. Finally, all subjects’ trials dFC were concatenated along the temporal dimension. We defined this matrix as a ‘Group dynamic functional connectivity matrix (G-dFC)’, denoted ‘X’.

#### 2.4. Task-related functional brain networks

##### Problem statement

The resultant G-dFC matrix representing the time-varying features can be expressed as as a linear mixture of elementary brain networks that fluctuate dynamically over time. Such issue is the main concern of Source Separation (SS) approach aiming at recovering ‘k’ hidden sources from a set of observations with minimal priori knowledge about these sources. In this context, the SS problem can be formulated as follows:

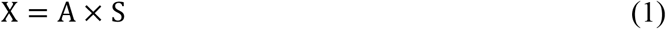

Where:

- ‘X’ is the computed G-dFC matrix of dimension [qxm]:

- q= Nx(N-1)/2 with N=78, representing connectivities between all ROIs.
- m=TxN_tot_ with T is the number of time windows and N_tot_ is the total number of trials for all subjects.
- ‘A’ is the mixing matrix of dimension [qxk] illustrating the contribution weights of each individual connection to the components sources, thus the spatial maps of brain networks (k<min(q,m)).
- ‘S’ is the sources matrix of dimension [kxm] representing temporal sources signatures of G-dFC, collapsed across all connections.

Although it is difficult to determine an optimal setting for the number of reduced components, ‘k’ was set to 10 in this study for all SS algorithms, consistent with several previous works. It was chosen as a trade-off value to be able to detect sufficient task-related sources without missing genuine components at the same time. The effect of k was evaluated in simulated data (see supplementary material). Therefore, our main objective is to uniformly explore several SS methods abilities to find the appropriate mixing and sources matrices of G-dFC. Among existing SS algorithms, we chose nine popular/well-known methods: six different variants of temporal Independent Component Analysis (tICA) (JADE, INFOMAX, SOBI, FastICA, COM2, and PSAUD), Principal Component Analysis (PCA), Non-negative Matrix Factorization (NMF) and Kmeans as a state-of-the-art clustering method. They all transform the desired matrix factorization into spatial maps and time series. However, they differ primarily in the constraints imposed on decomposed components. Below, we will give an succinct description about these methods.

##### Independent Component Analysis: ‘Temporal Statistical Independence’

ICA tends to linearly transform multivariate observations into a set of ‘statistically mutually independent’ latent variables under the hypothesis that these variables are as ‘non-Gaussian’ as possible. In our study, we examine temporal ICA (tICA) adopted by several previous studies(O’Neill et al., 2017; Yaesoubi et al., 2015) in order to obtain states that fluctuate independently in time. In this context, decomposed signals ‘S’ consist of the ‘k’ source time courses and the associated mixing matrix ‘A’ illustrates the contribution of temporally independent maps.

There are several criteria to measure independence such as minimization of mutual information and maximization of non-Gaussianity. Hence, different algorithms are proposed to perform ICA decomposition, each yielding to different ICA model with specific characteristics. Here, we evaluate tICA using six different popular and prominent methods: (1) JADE, (2) InfoMax, (3) SOBI, (4) FastICA, (5) CoM2 and (6) PSAUD. These methods are chosen in such a way to cover various statistical independence definitions, statistical order and computational process techniques. Briefly, InfoMax and FastICA are based on information theory, while all other selected methods optimize contrast functions based on cumulants of the data. Among them, SOBI uses only Second Order (SO) cumulants in contrast to others that exploit both SO and Fourth Order (FO) cumulants. In addition, FastICA and PSAUD use a deflation process for decomposition while other ICA variants jointly separate sources.

All ICA algorithms start with preprocessing steps including centering, whitening, and dimensionality reduction. Then, each method performs the source separation following its own criteria and hypothesis as described in supplementary materials.

##### Principal Component Analysis: ‘Variance maximization’

PCA is a basic linear technique widely used for data dimensionality reduction. It involves a mathematical procedure that transforms a set of observations of possibly correlated variables into smaller number of orthogonal, hence linearly uncorrelated variables called principal components or ‘eigenvectors. This procedure is defined in such a way that the variance or ‘eigenvalues’ of the data is maximized. Then, a fixed number ‘k’ of eigenvectors and their respective eigenvalues can be chosen to obtain a consistent representation of the data. Here, we apply the Singular Value Decomposition (SVD) algorithm of PCA(Golub and Reinsch, n.d.) on our predefined input matrix ‘X’, where eigenvectors and eigenvalues are computed from the correlation matrix of ‘X’ defined as:

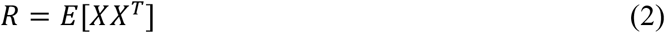

In this way,

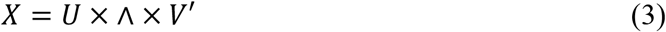

Where, ‘Λ’ is a diagonal matrix of the same dimension of ‘X’ with non-negative diagonal elements in decreasing order of variance, ‘U’ is a unitary matrix containing orthonormal eigenvectors and ‘V’ the corresponding temporal coefficients. Thus, the mixing matrix ‘A’ is computed as the first ‘k’ elements of *U* × Λ, and ‘S’ as the first ‘k’ elements of ‘V’’ coefficients.

##### Non-Negative Matrix Factorization: ‘Positivity’

Nonnegative matrix factorization (NMF) is an unsupervised machine-learning technique(Lee and Seung, 1999) that imposes ‘non-negativity’ constraint on the decomposed factors when solving SS problem. When applied to G-dFC data ‘X’, NMF leads to parts-based representation that captures additive combination of basis subgraphs ‘A’ at each time window with temporal coefficients ‘S’ eliminating negative signal variations. Among several existing NMF approaches, we selected Alternating Least Squares (ALS) algorithm that has previously shown good performance in fMRI context(Ding et al., 2013).

##### Kmeans Clustering: ‘Sparsity’

Kmeans is one of the simplest unsupervised learning algorithms that solve the SS problem through clustering approach(Lloyd, 1982). The algorithm works iteratively to assign each point to only one of the ‘k’ groups based on feature similarity. Mathematically, by considering G-dFC spatial connectivity as a set of m-dimensional vectors {C_1_, C_2_, …, C_m_}, the Kmeans clustering aims to generate a partition into ‘k’ clusters A = {A_1_, A_2_, …, A_k_} to minimize the sum of within-cluster distances ‘J’ over multiple iterations J:

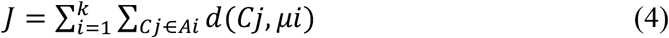

where μi is the mean of spatial connectivity in Ai, and d represents the distance between the two vectors. In our framework, the sparse coding adopted by Kmeans restricts a single time point to have a unique activated network state. The computed clusters ‘A’ represents the structure of common connectivity patterns across subjects. Then, ‘S’ is calculated as the frequency of reoccurring of each cluster at each time point through clusters indices. Here, we adapted the same procedure of Kmeans used by Allen et al.(Allen et al., 2014): L1 (Manhattan) distance is used, as it was suggested to be more effective than L2 (Euclidiean) distance for high-dimensional data(Aggarwal et al., 2001). The algorithm is replicated 100 times to increase chances of escaping local minima, and centroid positions were randomly initialized.

#### 2.5. Comparative Analysis

##### MEG group-level analysis

For each dataset, the SS method generated k=10 components. Among these 10 components, identifying those that reflect genuine brain activity related to the task is critical. In this paper, we followed a testing procedure adopted by(O’Neill et al., 2017) and previously described in(Hunt et al., 2012; Winkler et al., 2014) to determine significant components modulated by the tasks. The testing relies on the construction of empirical null distribution based on a ‘sign flipping’ permutation approach. In this context, we selected randomly half of the subjects and flipped the sign of their contribution to the extracted components. To give all possible realizations, each time a different set of subjects was selected for sign flipping, yielding to 6435 combinations for the self-paced data and 92378 for the Sternberg data. Regarding HCP Motor data, the same combination procedure could not be conducted directly on the total number of subjects for computational reasons. Therefore, we split the 61 subjects into 20, 20 and 21 subjects and performed separately three combinations, two consisting of 184756 repetitions and one of 352716 repetitions. In this way, the null distribution is built and an independent component was considered significant if, at any time point, the corresponding time signal, averaged over trials, fell outside a threshold defined at 0.05 with corrections. For all datasets, 2-tailed distribution was allowed, and Bonferroni corrections were applied for multiple comparisons across the 10 components and across temporal degree of freedom. It is assumed that a single temporal degree of freedom was added each time the window shifts by more than half of its width. This meant a total of 8 temporal degrees of freedom in the self-paced data, 12 in the Sternberg data and 6 in the HCP Motor data. Thresholds were therefore set at (0.05/(2*10*8)= 0.0003) for the self-paced experiment, (0.05/(2*10*12)= 0.0002) for Sternberg experiment and (0.05/(2*10*6)= 0.0004) for HCP Motor data.

##### MEG subject-level analysis

Besides group-level analysis, it is crucial to test the performance of each method when applied directly on individual dFC. To this end, instead of concatenating trials from all subjects as in the final step of ‘G-dFC’ computation, we perform, for each subject, a dFC concatenation of all trials related only to this subject to form a subject specific dFC, denoted ‘S-dFC’. Then, all selected SS methods were applied on ‘S-dFC’ matrix to extract subject-specific spatial and temporal signatures (k=10). In order to quantitatively evaluate and compare methods strength at subject-level context, we measure, for each method, both spatial and temporal similarities between each extracted S-dFC component and significant G-dFC components. These parameters are:

1. Average Distance (AD) for ‘spatial similarity’:

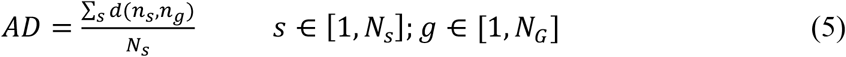 Where *d*(*n_s_,n_g_*) is the Euclidian distance between the node *n_s_* of S-dFC network and the nearest node *n_g_* from the significant G-dFC network. *N_s_* is the total number of nodes in S-dFC network, and *N_G_* denotes the total number of nodes in G-dFC network. All networks were 70% thresholded. Lower values of AD indicate stronger spatial similarity between S-dFC and G-dFC networks.
2. Correlation Signals (CS) for ‘Temporal Similarity’:

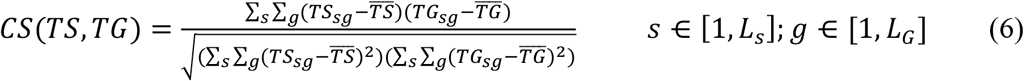 Where *TS* is the temporal signal of each S-dFC component of length *L_s_* and *TG* represents temporal signals of G-dFC significant component of length *L_G_*. Higher values of CS reveal stronger temporal similarity between S-dFC and G-dFC signals.

We perform this analysis on each subject among the 15 subjects of the MEG dataset 1 (Motor task). Therefore, for each method, we counted the number of subjects that show satisfactory results performance in the context of S-dFC, based on the previously explained measures. Then, to approximate the number of subjects/trials needed for each SS method to give significant results, we follow the same procedure explained above, but instead of single subject S-dFC computation, we increased the number of concatenated subjects in dFC computation from N_subj_=2 to 14, progressively. In order to have generalized and reliable results, we considered all possible combinations relative to each 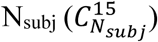, where different sets of N_subj_ subjects were selected among the 15 existing data subjects.

### 3. Data availability

Data supporting the findings of this study are available in the link (https://github.com/judytabbal/dynbrainSS.git). All HCP data are available on https://www.humanconnectome.org/software/hcp-meg-pipelines. The datasets 1 and 3 are available upon request.

### 4. Code availability

Codes supporting the findings of this study are available in the link (https://github.com/judytabbal/dynbrainSS.git). All analysis codes necessary to produce the results here were performed in MATLAB software, using FieldTrip Toolbox http://www.fieldtriptoolbox.org for data segmentation, filtering and source reconstruction steps, EEGLAB toolbox for some SS methods as JADE, InfoMax and SOBI and other MATLAB implemented functions as detailed and provided in the previous link. A graphical user interface is made freely available allowing other researchers to test the methods on our (or their) simulated and real data.

## Results

To track the fast reconfiguration of functional brain networks during cognition, we compared the performance of nine source separation (SS) methods, six variants of ICA (JADE, InfoMax, SOBI, FastICA, CoM2 and PSAUD), PCA, NMF and Kmeans. In the following, we present our evaluation study on real MEG data, however our methodology was also tested on simulated data. These results can be found in the supplementary material. Briefly, the simulation-based analysis showed that all methods provide satisfactory results in terms of spatial and temporal similarity between reconstructed and simulated components with the best performance for NMF method and the worst for SOBI. All methods, except for FastICA and Kmeans, provided consistent results. CoM2 and PCA were the fastest. Results demonstrated that FastICA, Kmeans and PSAUD depend on the imposed number of components while FastICA was affected by the noise level. Reader can refer to supplementary material to see the detailed quantitative analysis on simulated data.

The SS methods were compared on three MEG datasets consisting of two motor tasks i) self-paced (N=15), ii) HCP left hand movement (N=61) and iii) working memory task (N=19). Our goal was to extract the brain networks with the corresponding time course that are mainly involved during each task. To achieve this goal, subjects’ data were preprocessed and filtered and corresponding brain sources activity was reconstructed using Beamforming with reduction of spatial leakage in order to reduce the effects of volume conduction. Then, dynamic functional connectivity matrices were computed on time-windowed brain sources signals using Amplitude Envelope Correlation (AEC) measure. To obtain Group Dynamic Functional Connectivity (DFC), matrices were concatenated over all subjects and trials. We next compared the different SS methods on the average matrices obtained during the time-course of the distinct behavioral task.

Results are illustrated in Figures 2,3,4. In each figure, we presented only the components that demonstrated significant task modulation based on a null distribution-based approach (proposed in (O’Neill et al., 2017) and described in the methods section). The networks were thresholded only for visualization purpose. Corresponding dynamic reconfiguration of each significant network were plotted together. The temporal fluctuations represent component time signals averaged over trials and subjects.

**Figure 2.**
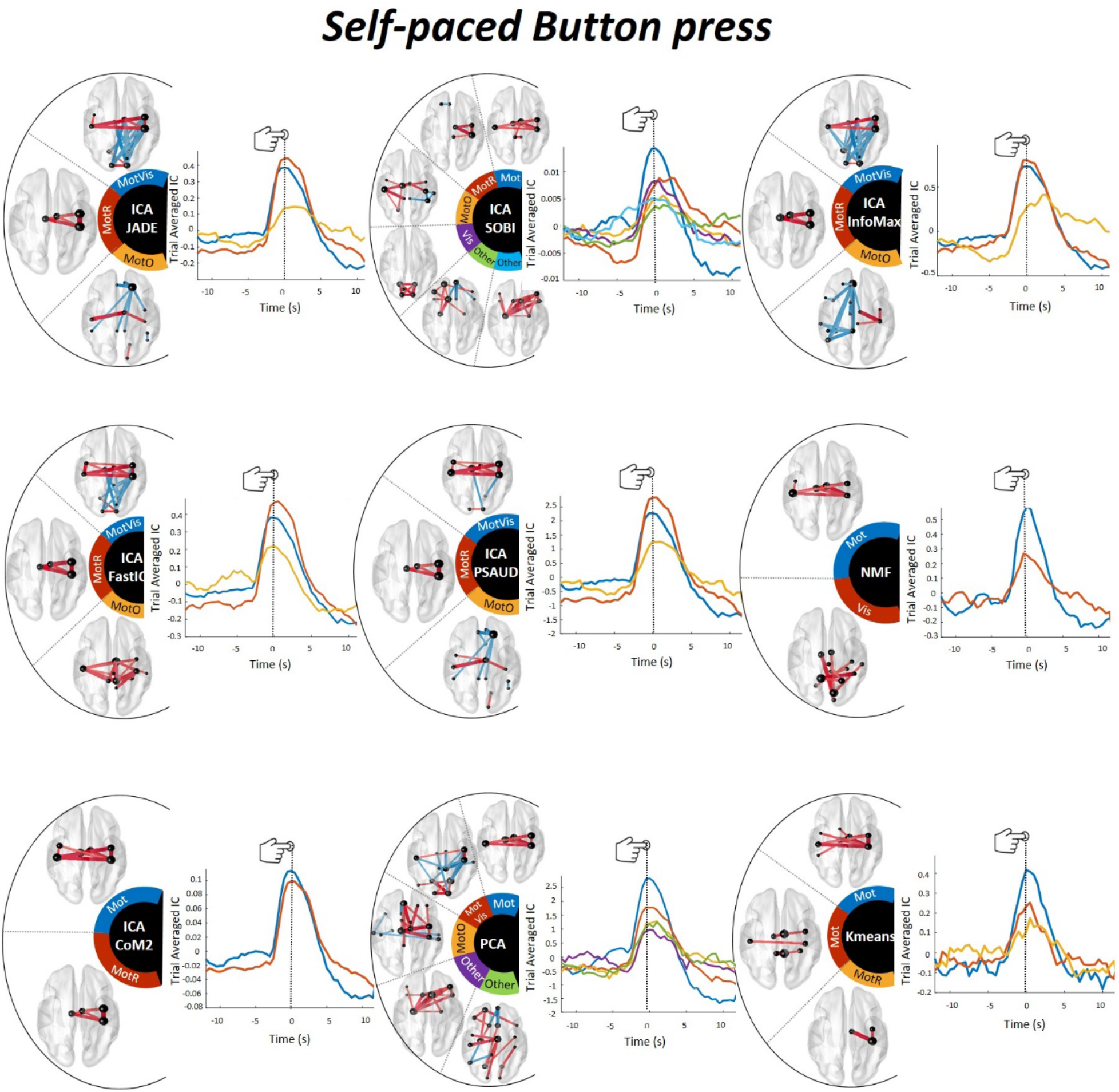
Self-paced motor task results. Spatial and temporal distribution of all significant components derived from all compared SS methods applied on G-dFC in the self-paced motor task (N=15 subjects). All brain networks were thresholded for visualization; lines width indicates connectivity strength between regions. Red lines represent positive connectivity values while the blue ones represent negative values. Integrated AAL nodes are represented by spheres of different sizes that reveal connectivity weights between that region and the rest of brain. Corresponding temporal evolution is averaged across all trials and subjects. Time values on the x-axis represent the position of the sliding window’s center, relative to the button press at t=0sec (as illustrated by a vertical line). A color code is attributed for each component in space and time. For each SS method, only significant components (pcorrected<0.05) that appear outside the ‘sign-flip’ based null distribution (as described in methods sections) are shown here. All 10 extracted components with corresponding null distribution are shown in Supplementary Figure S4 for an example of ICA-JADE method. Note that sensorimotor network is clearly activated at the button press instant in all SS methods. In this figure, ‘Mot’ refers to Motor network, ‘MotR’ to Right Motor, ‘Vis’ to Visual, ‘MotVis’ to Motor-Visual, ‘MotO’ to Motor-Others and ‘others’ to other networks. An interactive version of ICA-JADE results can be found on our github https://github.com/judytabbal/dynbrainSS.git using rotatable MATLAB figures.

**Figure 3.**
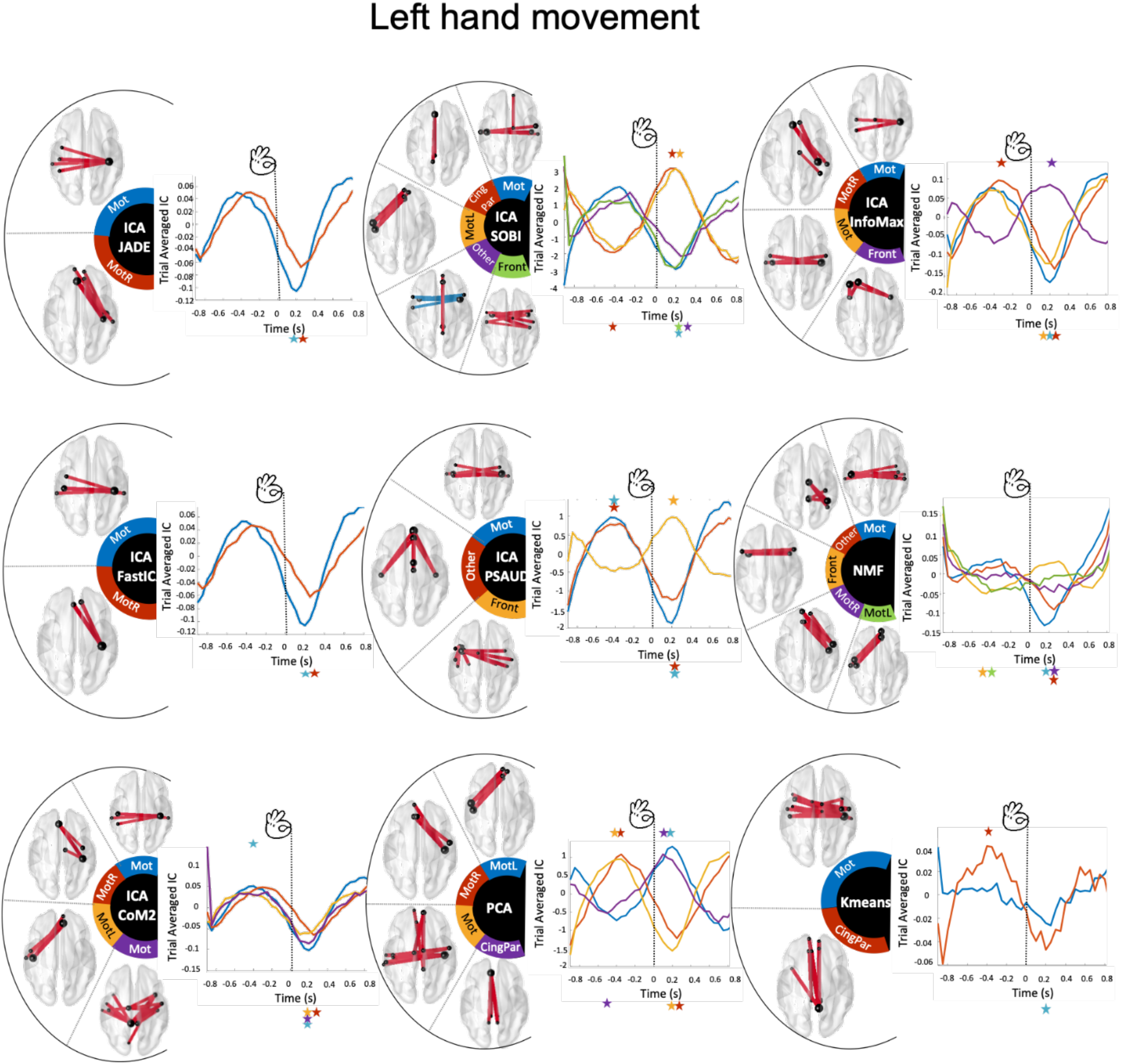
HCP left hand movements task results. Spatial and temporal distribution of all significant components derived from all compared SS methods applied on G-dFC in the lefthand motor task (N2=61subjects). All brain networks are thresholded for visualization. Time values on the x-axis represent the position of the sliding window’s center, relative to the button press at t=0sec (as illustrated by a vertical line). A color code is also attributed for each component in space and time. For each SS method, only significant components (pcorrected<0.05) that appear outside the null distribution are shown here. Exact times of significance relative to each component are indicated with stars., revealing an oscillatory temporal activation of motor component for all SS methods. All 10 extracted components with corresponding null distribution are shown in Supplementary Figure S5 for an example of ICA-JADE method. In this figure, ‘Mot’ refers to Motor network, ‘MotR’ to Right Motor, ‘MotL’ to Left Motor, ‘CingPar’ to Cingulate-Parietal, ‘Front’ to Frontal and ‘others’ to other networks. An interactive version of ICA-JADE results can be found on on our github https://github.com/judytabbal/dynbrainSS.git using rotatable MATLAB figures. Reproducing these results is also possible/available using the MATLAB interface on github.

**Figure 4.**
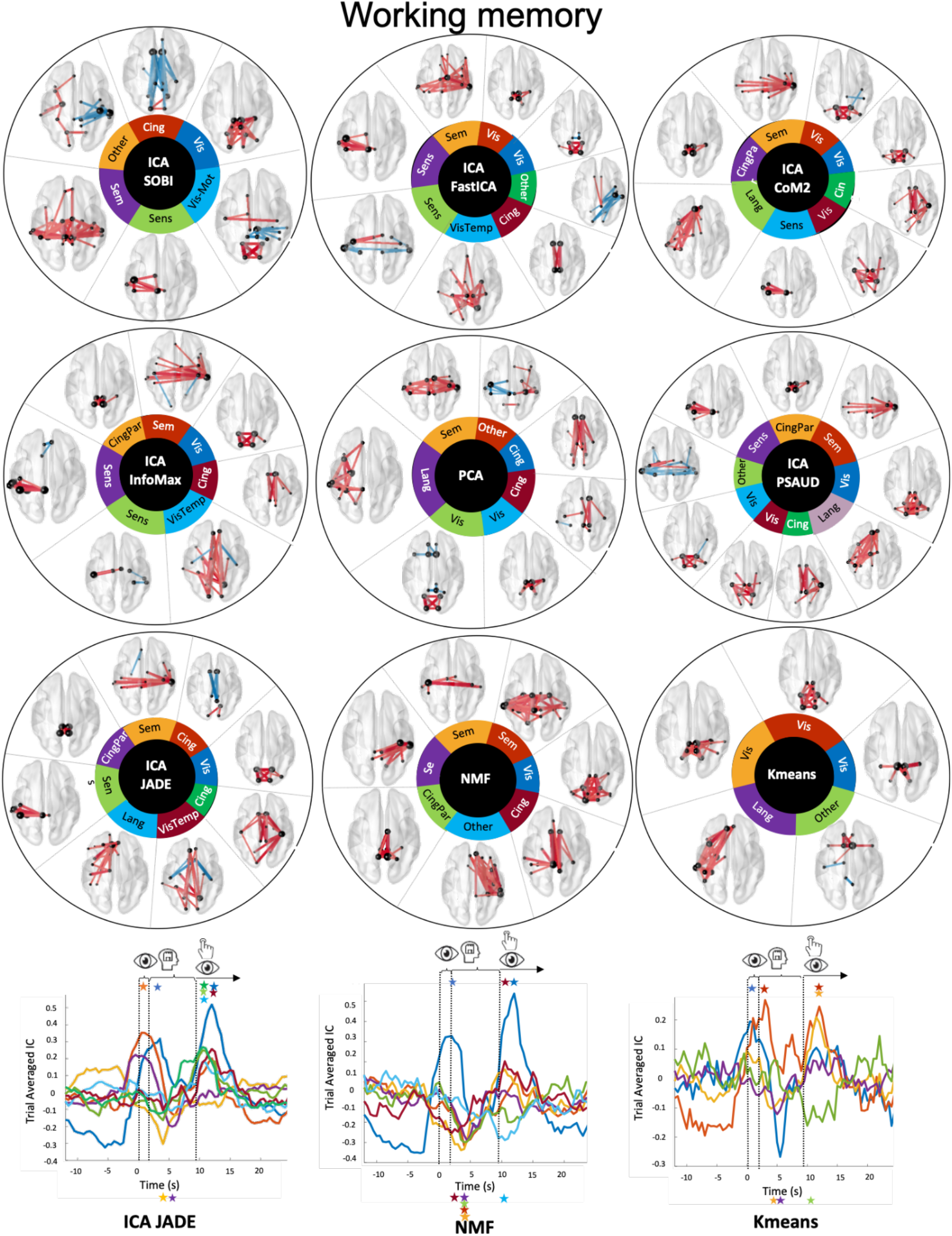
Sternberg working memory task results. Spatial and temporal distribution of all significant components derived from all compared SS methods applied on G-dFC in the working memory task (N_3_=19subjects). All brain networks are thresholded for visualization. Time values on the x-axis represent the position of the sliding window’s center, relative to the first visual stimulus presentation at t=0sec (as illustrated by a vertical line). The first two vertical lines illustrate the instant of successive visual examples presentation at t=0s and 1.6sec and the third vertical line at t~9sec separates between the maintenance period that lasts for 7sec and the probe presentation followed by a possible button press and feedback. A color code is attributed for each component in space and time. For each SS method, only significant components (p_corrected_<0.05) that appear outside the null distribution are shown here. Exact times of significance relative to each component are indicated with stars. Temporal variation of only JADE, NMF and Kmeans is illustrated, whereas the rest are shown in Supplementary Figure S7. All 10 extracted components with corresponding null distribution are shown in Supplementary Figure S6 for an example of ICA-JADE method. Note that in this task, much larger variety of significant networks are extracted among SS methods, including visual, sensorimotor, language, semantic, cingulate and visio-temporal network at different temporal activation. In this figure, ‘Vis’ refers to Visual network, ‘Sem’ to Semantic, ‘Sens’ to Sensorimotor, ‘Lang’ to Language, ‘CingPar’ to Cingulate-Parietal, ‘Cing’ to Cingulate, ‘VisTemp’ to Visual-Temporal and ‘others’ to other networks. An interactive version of ICA-JADE results can be found on our github https://github.com/judytabbal/dynbrainSS.git using rotatable MATLAB figures.

### Self-paced button press task

In this task, participants were asked to press a button with the index of their non-dominant hand every 30 seconds. Based on the results obtained by all SS methods, we divide all derived significant components into 6 networks, as denoted in Figure 2: Motor (‘Mot’), Right Motor (‘MotR’), Visual (‘Vis’), Motor-Visual (‘MotVis’), Motor-Other (‘MotO’) and ‘other’. Detailed description of brain regions integrated in each of these networks can be found in supplementary materials. The four ICA variants (JADE, InfoMax, FastICA and PSAUD) lead to very similar results where three significant networks were extracted: ‘MotVis’, ‘MotR’ and ‘MotO’ networks. CoM2 and Kmeans show ‘Mot’ and ‘MotR’ networks, whereas ‘Mot’ and ‘Vis’ networks arise from NMF decomposition. Regarding SOBI and PCA, we can see higher number of components: ‘Mot’, ‘MotR’, ‘MotO’, ‘Vis’ and two ‘other’ networks with SOBI, and ‘Mot’, ‘MotVis’, ‘MotO’, and two ‘other’ networks with PCA. Temporal variation was similar for all significant components over all methods showing a peak value at 0sec, the button press time, with slight differences in amplitude values, indicating signal intensities relative to each component. Note that negative connectivity, referred to as blue connections in spatial networks and negative temporal values in temporal signals, represents desynchronization between brain regions.

Therefore, all studied SS methods were able to extract networks that highlight strong connections between sensory and motor regions modulated significantly by the task at the exact button press instant (‘Mot’). Also, visual network has shown significant modulation, either with motor connections (‘MotVis’) as in JADE, InfoMax, FastICA, PSAUD and PCA, or as a single component (‘Vis’) in SOBI and NMF. In addition, all methods except for CoM2, NMF and Kmeans, derived an additional significant network that links sensorimotor cortex to other brain regions (‘MotO’). For SOBI and PCA, further two ‘other’ components were revealed significant.

In summary, all methods were able to extract at least one significant motor network at button press time, while SOBI and PCA selected extra components, not directly related to the studied task, as significant components.

### Left-hand movement task

This task is also motor but different than the previous one. Here the participants were asked to rapidly and successively tap their left index and thumb fingers. Six network patterns were observed over all the methods, as denoted in Figure 3: Motor (‘Mot’), Right Motor (‘MotR’), Left motor (‘MotL’), Cingulate-parietal (‘CingPar’), Frontal (‘Front’) and ‘other’. Implicated brain regions are described for each network in supplementary material. Figure 3 shows that at least one ‘Mot’ network is extracted by all methods. This network shows significant drop in connectivity around 0.2sec following the button press instant. CoM2, PSAUD and PCA show also significant increased modulation of this network at −0.2sec. Exact times of components significance are indicated with stars on Figure 3. Another motor network ‘MotR’ was extracted from JADE, InfoMax, FastICA, CoM2, PCA and NMF methods. This network follows the same temporal variation of the previous one. Interestingly, this network emphasizes on the activation of right motor cortex that is consistent with the nature of the task (left hand movements). SOBI, CoM2, PCA and NMF methods have also extracted a ‘MotL’ network. This network seems to have opposite temporal variation from the previous ones (significant increased connectivity at +0.2sec for SOBI and PCA, and decreased connectivity at −0.2sec for NMF). Two networks ‘CingPar’ and ‘Front’ were commonly derived by SOBI, PCA and Kmeans for the former and InfoMax, SOBI, PSAUD and NMF for the latter. Both networks show decreased modulation around −0.4sec then an increase around 0.2sec.

The ‘other’ networks were found in SOBI (inferior frontal to temporal connections decorrelated with medial frontal and cingulate connections), PSAUD (medial frontal with temporal, motor and cingulate connections) and NMF (mid and post cingulate with parietal and motor connections) as illustrated in the Figure 3. Regarding temporal evolution, the hand movement here are much more frequent than the previous task. Clearly, the fast neural activity due to the short time between successive button presses is expressed as an oscillatory behaviour of brain network activity around the zero time button press, as showed also previously(Vidaurre et al., 2018a). This yields to the obtained temporal variation where the motor network state seems to have high connectivity before button press and begin to have a drop-in connectivity to reach its significant peak after ~0.2sec (referred to as a desynchronization in high frequencies(Vidaurre et al., 2018a)). In this context, the same oscillatory time variation is followed by other networks with similar or opposite modulation sign.

In summary, results show that all methods were able to extract a task-related component (‘Mot’) but some methods were more (with lower false positives) to extract the motor networks (JADE, FastICA and CoM2) than other methods.

### Working memory task

This task is much more complex comparing to the other two tasks. Subjects here were asked to visualize and memorize two visual shapes and respond to a third probe stimulus by a button press (with their right index finger) in case of matching. The increased cognitive load evoked by the Sternberg task is expected to induce variations in a greater number of significant brain networks including stimulus visualisation (visual network), semantic processing and pattern recognition (semantic, language networks) and button press response (sensorimotor network). Eight networks were obtained from the different methods: Visual (‘Vis’), Semantic (‘Sem’), Sensorimotor (‘Sens’), Language (‘Lang’), Cingulate-Parietal (‘CingPar’), Cingulate (‘Cing’), Visual-Temporal (‘VisTemp’), ‘other’. Reader can refer to supplementary material for brain regions description. Figure 4 illustrates the spatial distribution of all significant components for all SS methods, and temporal variation for three of these methods (JADE, NMF and Kmeans) for visualisation clarity. The temporal evolution of other SS methods can be found in Supplementary Figure S7. Starting from t=0sec, two visual stimulus (shapes) were presented successively, each for 0.6sec. During this period, all methods were able to extract one or more significant ‘Vis’ network. Some auxiliary connections to parietal or temporal regions can be seen in SOBI, FastICA2, PSAUD, NMF and Kmeans. Time variation of this network shows significant peak during the first two seconds period. At the same time, we can notice a decreased modulation in ‘Cing’ network connectivity in JADE, SOBI and PCA.

Following stimulus presentation, subjects should retain the observed shapes in working memory. During this period, known as the maintenance phase, all methods, except Kmeans, show significant decreased modulation at [4-6sec] of a ‘Sem’ network. The breakdown in this network’s connectivity was previously demonstrated(O’Neill et al., 2017). During the same period, we can notice a drop-in connectivity in ‘CingPar’ network as revealed by some methods (JADE, InfoMax, CoM2, PSAUD and NMF). At [10-12sec] period, many networks seem to be significantly modulated among all methods: (1) ‘Vis’ is re-activated at the probe stimulus presentation in all methods in the form of one (JADE, InfoMax, SOBI, FastICA and NMF), two (PSAUD, PCA and Kmeans) or three (CoM2) components. In these networks, we can notice some additional connections to parietal or temporal nodes as in SOBI, PCA, NMF, and Kmeans, or to parahippocampal gyrus as in CoM2 and PSAUD, (2) ‘Sens’ shows significant increase with all ICA subtypes methods. This network becomes most strongly connected around the time button press response, (3) ‘Lang’ is commonly derived by JADE, CoM2, PSAUD, PCA and Kmeans and exhibits an increased connectivity peaking during probe presentation. We can notice that this network is also significantly decreased in CoM2 and Kmeans around 5sec. Connections to motor and frontal node are observed in JADE, and to visual lingual in Kmeans, (4) ‘Cing’ in JADE, InfoMax, FastICA, CoM2, PSAUD, PCA and NMF. InfoMax, PCA and NMF have few links to parietal lobe and (5) ‘VisTemp’ network was obtained in JADE, InfoMax and FastICA. Besides these components, ‘other’ network was considered significant in many methods as SOBI, FastICA, PSAUD, PCA, NMF and Kmeans. These components are illustrated and denoted in the Figure 4.

In summary, the three ICA variants (JADE, CoM2 and PSAUD) succeeded to derive all expected components and show their appropriate temporal significant modulation. Other ICA variants (InfoMax, SOBI and FastICA) were able to extract visual, semantic and sensorimotor but not the language network. PCA missed the sensorimotor components. However, NMF was unable to show both sensorimotor and language networks and Kmeans failed to extract semantic and sensorimotor components. Overall, one can appreciate more the ICA results in this task, especially for JADE, CoM2 and PSAUD. NMF and Kmeans tend to extract task-related significant components but missing important networks. For the three tasks, spatial and temporal distribution of the 10 components derived from JADE method, with corresponding null distribution, can be found in Supplementary Figures S4, S5, S6. For further clarity in visualisation and interpretation, we illustrated, in Figure 5, the spatiotemporal reconfiguration of the functional brain networks as obtained by ICA-JADE.

**Figure 5.**
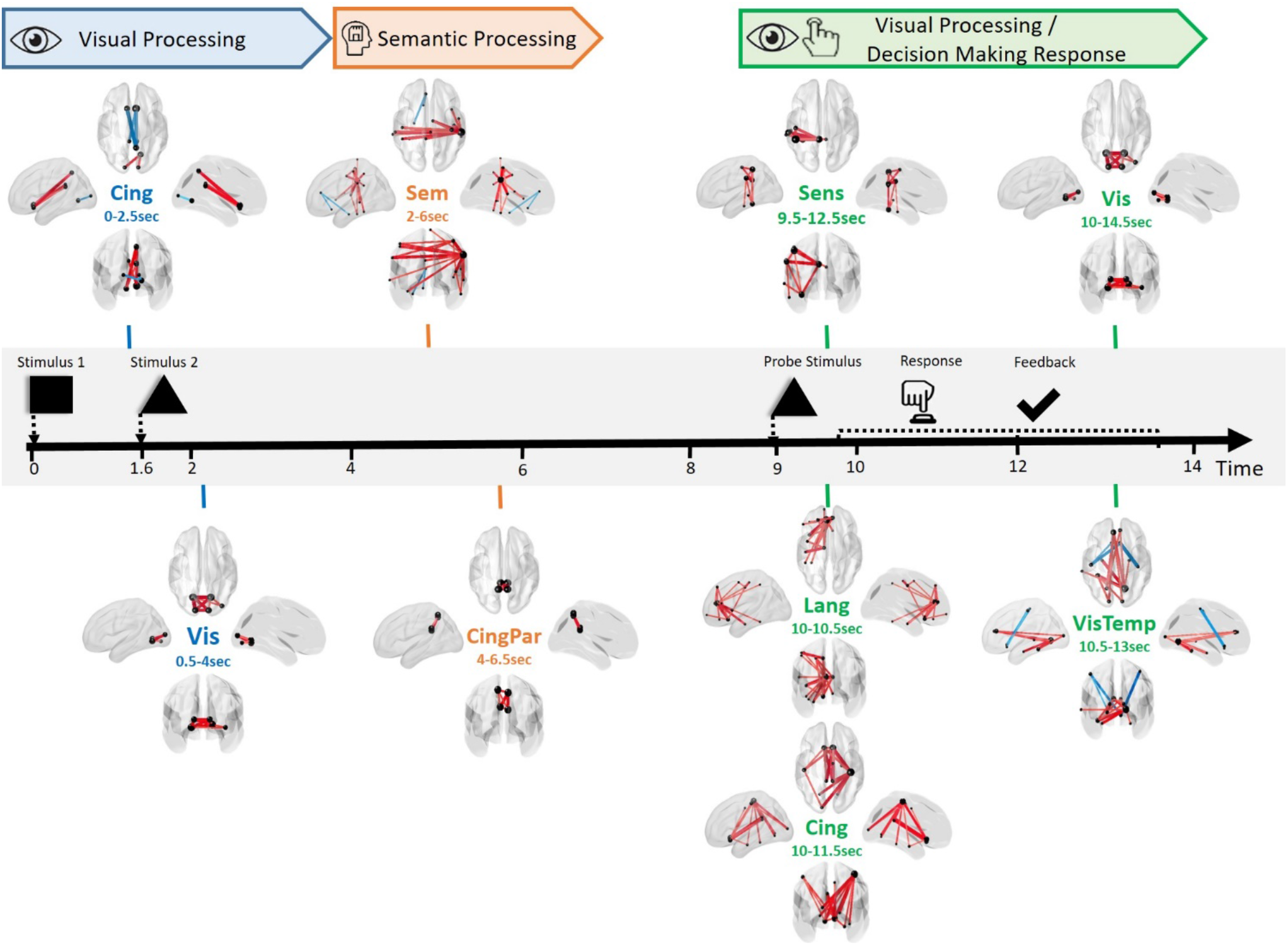
Typical example of the spatiotemporal reconfiguration of brain networks during working memory task using ICA-JADE. All significant networks extracted from JADE are collected and presented sequentially relative to each event and period time. The nomination and the exact temporal period of significant activation of each network is clearly indicated. Corresponding cognitive functions are also specified. In this figure, ‘Vis’ refers to Visual network, ‘Sem’ to Semantic, ‘Sens’ to Sensorimotor, ‘Lang’ to Language, ‘CingPar’ to Cingulate-Parietal, ‘Cing’ to Cingulate and ‘VisTemp’ to Visual-Temporal.

Furthermore, we discriminated the different SS methods performance in terms of activated brain networks proportion in each task. For example, in motor tasks, we compared the proportion (referred as ‘weight’) of all extracted motor networks (Mot, MotR, MotVis, MotO) with that of auxiliary networks (that did not involve motor regions). For working memory task, visual, semantic, language, sensorimotor and auxiliary networks were evaluated with all remaining significant components occupancies for each method. Results are shown in Figure 6 with the spatial representation of the corresponding relevant brain regions.

**Figure 6.**
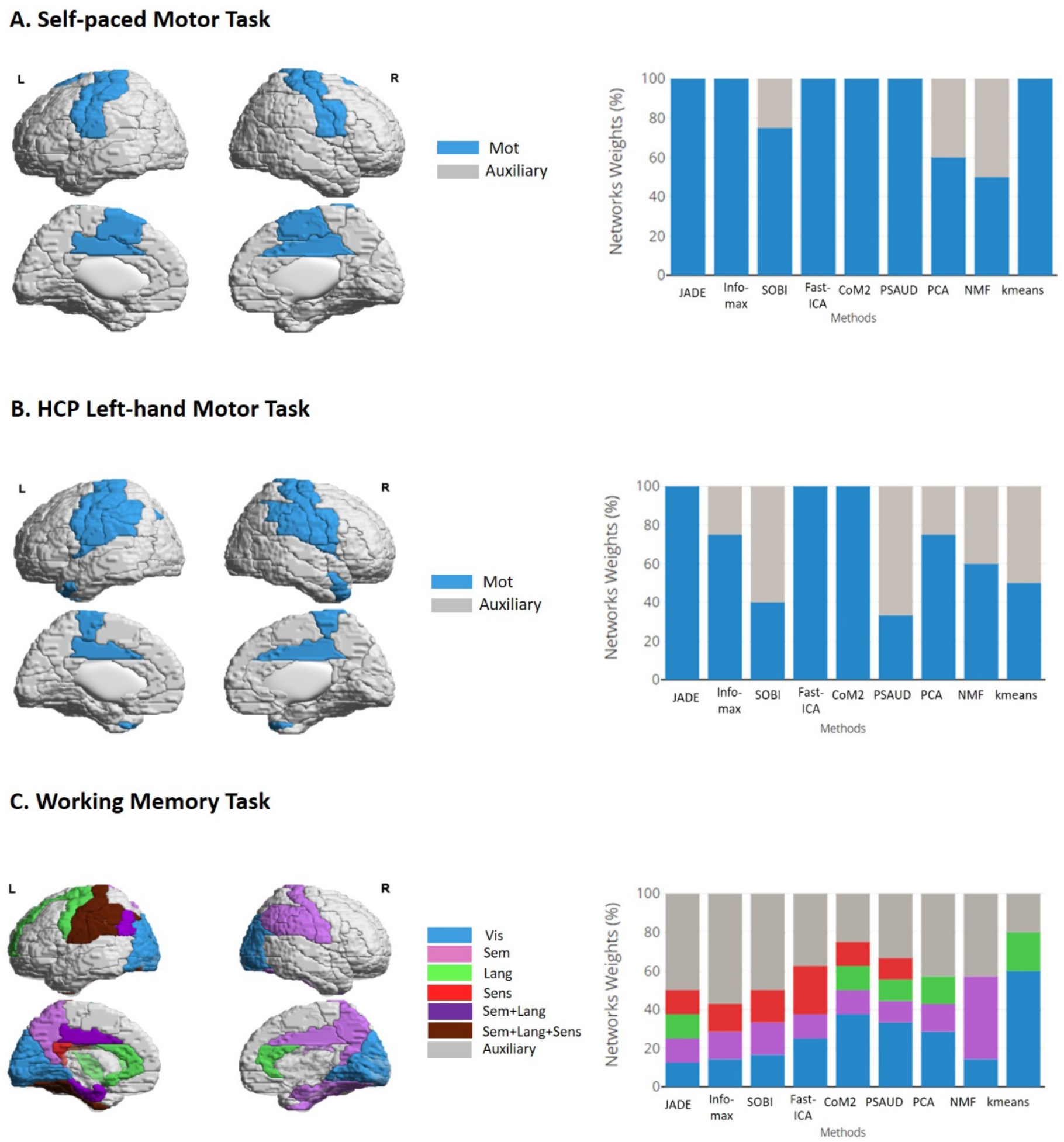
SS methods performance evaluation for real MEG tasks. For each task, brain regions involved in each relevant network are illustrated on the left side while brain networks weights are shown on the right side. The weights express the percentage occupancy of a defined brain network relative to all significant extracted components. For motor tasks, motor network (‘Mot’) was emphasized with auxiliary networks relative to all significant components, while visual (‘Vis’), semantic (‘Sem’), language (‘Lang’), sensorimotor (‘Sens’) networks are highlighted in contrast to other auxiliary networks. Referring to these representations, capabilities of different SS methods in extracting task-related components can be evaluated.

### Performance of each SS methods at subject-level

Here our objective is to evaluate the performance of the methods at the subject-level. We test i) the capacity of each method to extract significant components related to the task: to do so, we computed the correlation between the components obtained by each method on each subject with the significant network obtained at the group level and ii) the number of subjects needed for each method to detect ‘expected’ networks: here we tested the overall performance of each method by increasing the number of subjects, going from 1 to 15 as we performed subject-level analysis on the self-paced data. Figure 7.A summarizes the subject-level analysis scenario. For each method, we chose one of the significant motor components derived from the decomposition of the group-level (N=15subjects), mostly the one showing little intervention from regions other than sensorimotor (‘Mot’ or ‘MotR’) and having high temporal coefficients amplitude (supposed to be the best for each method). This component illustrates a ‘group’ motor network with temporal modulation at the button press time, and will, eventually, serve as a ‘mask’ component for subjectlevel analysis.

**Figure 7.**
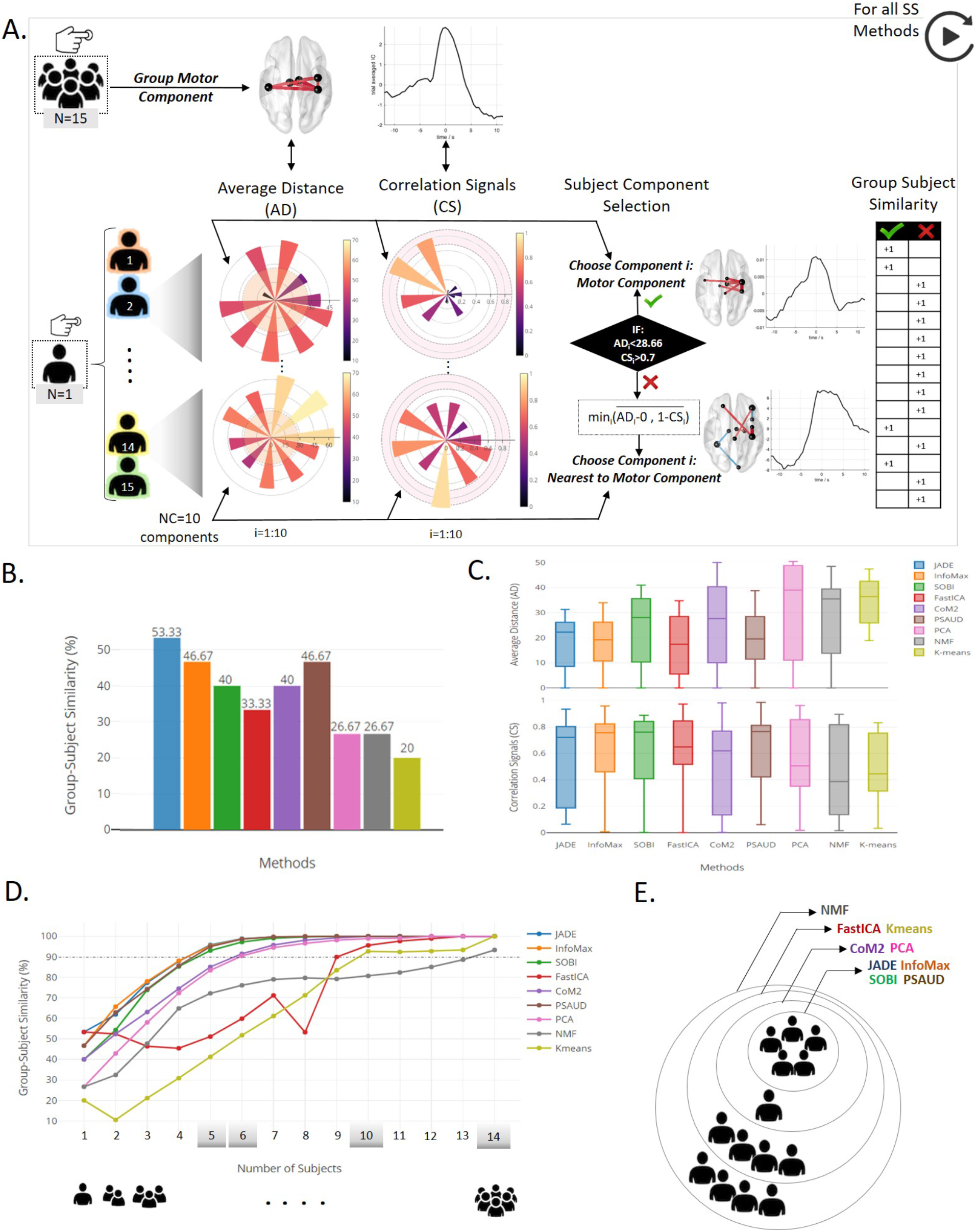
Subject-level analysis and results relative to the self-paced motor experiment. A. description of subject component selection procedure based on Average Distance (AD) and Correlation Signals (CS) values between subject’s component and the group motor component, when SS methods are applied on S-dFC data. Based on AD and CS condition limits, success and failure in extracting motor component cases are both illustrated by subjects 2 and 14 respectively. Spatial and temporal distribution with corresponding AD and CS values for all 10 components of both subjects 2 and 14 are illustrated in Supplementary Figure S8. B. Results of the number of subjects that successfully extracted a motor component in each SS method relative to the total subject’s number, denoted Group-Subject Similarity, are shown. C. distributions of AD and CS values of all selected components for the 15 subjects are illustrated. D. Generalization study with increasing number of subjects and the corresponding results of Group-Subject similarity percentage (when considering all possible combinations). E. Grey shaded values (5,6,10 and 14) represent the critical number of subjects required for each SS method to have a 90% precision in extracting the task related component.

For each SS method, NC=10 components were derived from each subject data. Then, Average Distance (AD) and Correlation Signals (CS) between each of these 10 components and the ‘group’ motor component relative to the SS method were computed in order to quantify the ability of the method to extract, from a single subject, a task-related component in space (motor network) and time (temporal modulation at button press time) respectively. Following this, only one of these 10 components is selected for results calculation. This selection is based on two conditions criteria on AD and CS values. In the case where AD component is less than a threshold (set as the average of AD values of all components for all subjects and SS methods), and CS is higher than 0.7, then the component is considered to be a motor component. Thus, if at least one of the 10 components pass these conditions, corresponding AD and CS values are denoted and the number of subjects that give similar results to group-level is raised by 1. Otherwise, we selected component *i* as the nearest component to the group-level result, with a compromise between spatial and temporal similarities.

A typical example is illustrated in Figure 7.A showing that PCA decomposition was able to extract a motor component from subject 2 (component 5), whereas no motor component was derived from subject 14, AD and CS values of component 7 were denoted in this case. Spatial and temporal distributions of selected subjects’ components in both cases are shown in Figure 7. Results of the other 9 components for these 2 subjects’ examples are shown in Supplementary Figure S8.

As a result, two parameters were collected and represented in Figure 7.B and 7.C respectively. Group-subject similarity percentage was calculated as the number of subjects that gives a motor component similar to the group-level result relative to the total number of subjects (N=15). Figure 7.B illustrates this parameter for all SS methods. We can see that JADE was able to extract a task-related component from 8 out of 15 subjects (53.33%), InfoMax and PSAUD from 7 subjects (46.67%), SOBI and CoM2 from 6 (40%), FastICA from 5 (33%), PCA and NMF from 4 (26.67%) and Kmeans from 3 (20%). The Figure 7.C shows the distributions of AD and CS values of selected components from each subject over all SS methods. Methods with higher subject-group similarity percentage have lower median values of AD and higher median values of CS. In addition, we can notice from AD and CS median values that similarity in space was much easier to be satisfied than temporal similarity for most SS methods. Interquartile range values show the existence of intersubject variability results. However, some methods showed higher interquartile range of AD values (CoM2 and PCA), or CS values (JADE, CoM2 and NMF) relative to other methods.

### The optimal number of subjects of each SS method

Then, the same procedure was applied with increasing the number of subjects from one subject (single-subject) to 14 subjects. AD and CS are computed for all possible combinations. The number of possible combinations is calculated. For example, 7-subjects analysis requires 6435 combinations, hence 6435 values of AD and CS. For each number of subjects, we calculated subject to group similarity as the ratio between number of combinations that succeeded in extracting a motor component relative to the total number of possible combinations. Results are illustrated in Figure 7.E, F. As expected, the percentage similarity increases with increasing number of subjects for all SS methods. Fluctuations in similarity results are observed in some methods (as FastICA, NMF and Kmeans) due to the non-consistency characteristic of these methods (as previously proved). Some methods required a smaller number of subjects for data analysis to provide satisfactory results (motor component at button press time in our case) than others. For example, the four ICA versions (JADE, InfoMax, SOBI and PSAUD) required 5 subjects in order to attain a minimum similarity level of 90% between subject and group level results. CoM2 and PCA required 6 subjects, while much more subjects were needed for others (10 subjects for FastICA and Kmeans and 14 for NMF) as showed in Figure 7.E. Overall, ICA subtypes work better on subject-level data than other SS methods.

Overall, ICA methods and specially those based on the high order statistics (such as JADE), outperform other methods in extracting networks at the subject-level.

## Discussion

Emerging evidence show that discrete and repetitive brain states govern the spatio-temporal dynamics of functional networks. The identification of these connectivity patterns can be performed using different SS methods where each one imposes its specific constraints which yield to different results. Therefore, recent studies have been conducted to evaluate and compare the performance of SS methods in describing the brain states fluctuating over time(Leonardi, n.d.; Meyer-Baese et al., 2004; Miller et al., 2016; Xie et al., 2017). However, these studies suffer at least from one of these limitations: (1) applying a stationarity framework-based comparison, (2) considering a limited number of compared SS methods, (2) using only one database and (3) lack of simulation-based comparison providing a ground-truth validation.

In this study, we have systematically evaluated the robustness of the most popular SS methods applied to extract the main brain networks fluctuating during time in order to help researchers make a rational choice among the multitude of available methods. Specifically, nine algorithms have been compared using simulated data (see supplementary materials) and three independent MEG datasets (N=95) recorded during motor and memory tasks. The discrepancy in the datasets size and behavioral tasks performed allows testing SS methods performance on different scenarios. As the evoked responses (analyzed here) last for hundreds of milliseconds, we conducted our comparative analysis based on MEG datasets to benefit from the excellent temporal resolution of this technique. However, the same pipeline study can be applied in task-related fMRI context.

Our results show mainly that temporal ICA methods reached the highest spatial and temporal accuracies compared to other SS methods. Among ICA subtypes, high statistical order-based methods induce more robust results than those based on lower order. Therefore, brain components are better described as non-Gaussian independent sources. When considering multiple criteria, JADE and CoM2 methods outperform other SS methods, as we will discuss later in details.

The quantitative comparison performed on simulated dynamic networks showed that all SS methods have successfully separated functional networks based on their connectivity time courses. However, spatial and temporal similarities in SOBI were significantly lower than other SS methods, especially for the second simulated state (P2) as shown in Supplementary Figure S3, which involves more complex spatiotemporal activity than other states. As expected, FastICA, NMF and Kmeans methods were proved inconsistent with multiple runs. This is caused by the nature of these algorithms that is based on random input initialisations until solution convergence. Thus, it is important to keep in mind these limitations especially for clinical related studies, where results consistency is of high interest. In addition, results revealed that the performance of FastICA, PSAUD and Kmeans depend on the selection of the number of components. This is an important property to take into account when dealing with complex cognitive tasks where the number of involved brain states is hard to predict. The noise effect on the results obtained was also tested and showed that FastICA seems to be the only method affected by the noise level of data. Hence, FastICA is less recommended to be applied when working on electrophysiological data that appears to be noisy. Regarding computation time, CoM2, PCA and PSAUD were the fastest whereas InfoMax and JADE were the slowest. Still, the executed time of these algorithms is sensible to dataset’s features (size, complexity, type…), and may therefore vary between different datasets. However, it is noteworthy to mention that this criterion is not enough to determine the computational complexity of each method; other metrics such as the number of floating-point operations (FLOPs) required for the algorithm completion could be also tested. We also suggest for future studies to explore other data simulation approaches that build the desired ground-truth brain states based on more realistic modeling.

Methods performance were evaluated on three real MEG datasets already published and tested by previous studies(Casorso et al., 2019; O’Neill et al., 2017; Tewarie et al., 2019; Vidaurre et al., 2018a; Zhu et al., 2020). According to the motor tasks (self-paced and HCP left hand movements tasks), results clearly showed that all SS methods have successfully extracted one or more significant networks that involve strong connectivity between sensorimotor regions (‘Mot’) for both tasks. These regions mainly include nodes from central gyrus in the case of self-paced task, and central gyrus in the case of HCP task. The implicated networks are strongly coherent with the task(Melnik et al., 2017; Yousry, 1997) since it requires both movement (through button press or hand movement) and tactile response (as the subject will feel the button or his fingers tape). The effect of right-handedness of all participants of self-paced dataset is also revealed by the presence of motor network, where only sensorimotor regions from the right cortex are implicated (‘MotR’). Yet, this network is not present in PCA and NMF methods. Similarly, ‘left’ hand integration is revealed by the presence of ‘MotR’ in most SS methods in HCP dataset. Note that all methods, except CoM2 and Kmeans, highlighted significant connections in the visual lobe (Figure 2) as previously noticed by Oneill et al. studying the same task (O’Neill et al., 2017). This can be interpreted as a cross modal synchronization between visual and sensorimotor cortex as previously studied(Bauer et al., 2020). We can also explain visual nodes activation by the fact that subjects may try to imagine the ‘self’ cue to press the button at each trial, similarly to the motor imagery context. Regarding temporal evolution, it is clear that all networks modulate significantly with the exact button press time for self-paced task. Differently, the temporal variation related to HCP motor task takes an oscillatory shape, which was also reported by other studies dealing with the same dataset(Vidaurre et al., 2018a; Zhu et al., 2020). A possible reason for this activity was suggested by Vidaurre et al.(Vidaurre et al., 2018a) considering a leakage effect of temporal activity of previous button press into the next trial due to the fast successive trials. In both tasks, SOBI and PCA extracted components not directly related to the motor cortex. One possible explanation of these methods’ fragility is that they are both based on second order cumulant in separating sources. In addition, HCP results revealed the superiority of JADE, FastICA and CoM2 over other methods as they better reflect integrity for motor task related networks.

In order to evaluate the spatial and temporal accuracy of SS methods at higher levels of complexity, we tested the methods on Sternberg working memory task. All SS methods strongly highlighted visual network, which is consistent with the presentation of visual stimuli at two different times. Regions in the primary visual lobe related to stimulus visualisation(Grill-Spector et al., 1998) and lateral occipital cortex responsible for object/shape recognition(Corbetta et al., 1991; Grill-Spector et al., 2001; Kourtzi and Kanwisher, 2001) were present in these networks. The button press response is reflected by sensorimotor connections only using ICA subtypes methods consistently with previous working memory studies(Metzak et al., 2011; Yamashita et al., 2015). Strong integration of left cortex in this network may be explained by the fact that right-handed participants were asked to use their right index finger to press the button in case of matching stimuli. In order to process and maintain observed stimuli as a way to memorize them, a higher level of cognition is illustrated by a ‘semantic network’, which mainly encompasses bilateral parietal and temporal areas activation in all SS methods, except for Kmeans. This is coherent with previous studies that demonstrate the evident role of parietal cortex as a workspace for sensory and perceptual processing in working memory framework (Chai et al., 2018) through angular(Frackowiak, 1992; Vandenberghe et al., 1996), precuneus(Cavanna and Trimble, 2006), and hippocampal(Baddeley et al., 2011) areas. Bilateral inferior temporal regions also play important role in semantic processing(Nestor et al., 2006; Vigneau et al., 2006). Fusiform gyri, strongly modulated in our results, has also shown a particular concern in this context(Mion et al., 2010). The extraction of the ‘language’ network by JADE, CoM2, PSAUD, PCA and Kmeans methods, was compatible with previous findings(Brookes et al., 2011b; O’Neill et al., 2017). This left lateralised network includes connections from inferior frontal gyrus (more robust in JADE method) to nodes spanning central lobe (as in JADE and CoM2). Temporal and parietal lobes were remarkably activated by these methods, mainly the parahippocampal and supramarginal gyri respectively. These regions are critical in memory encoding and retrieval and semantic cognition(Axmacher et al., 2008; Caminiti et al., 2015; Demb et al., 1995; Derrfuss et al., 2004; Deschamps et al., 2014; Vigneau et al., 2006). In a similar (abstract shape based) working memory task, the interpretation of this network was related to a verbalisation naming strategy employed by participants as a way to aid in memory encoding(Caminiti et al., 2015; O’Neill et al., 2017). Therefore, this network’s activation may be possible with the task as it modulates strongly with the probe presentation and response time.

Although the main objective of this study is to help researchers choosing the appropriate SS methods in their studies, the applications of such dFC analysis/framework in the clinical domain is such a promising topic. Many aspects of motor abilities and cognitive competences, which can be revealed by motor and working memory tasks respectively, are demonstrated to be perturbed for patients. Consequently, future studies can apply the methodology used in the present paper in order to extract spatiotemporal dynamics of networks from both control and patients groups for a better understanding of dysconnectivity induced by neurological disorders(van den Heuvel and Hulshoff Pol, 2010). We can go beyond this application towards establishing a ‘neuromarker’ for clinical diagnosis. This requires the analysis of subject-specific dynamic connectivity states rather than examining differences in dynamic network properties. To this end, some previous studies worked on the assessment of subject-specific brain states using results of group-level analysis through back-reconstruction approach such as group information guided ICA (GIG-ICA)(Du and Fan, 2013) or spatio-temporal (dual)-regression(Beckmann et al., 2009). However, we argue that these results may depend on the quality and quantity of group data. Alternatively, in order to satisfy clinical demands and ensure consistent and reliable results on single subject data, we tested SS methods performance when directly applied on subject-level dFC (S-dFC) as previously explained. Results of our analysis revealed the superiority of ICA-based approaches, as they tend to extract task related component with less required number of subjects/trials than other SS methods. One explanation is related to the algorithm dependency on input data dimensions. For example, it is known that clustering methods as Kmeans require large set of informative input to adequately cluster sources(Dolnicar, n.d.). On the other hand, less number of data points are necessary for ICA algorithms. This question remains challenging due to the inter-subject variability as revealed by the timecourses of connectivity in the constructed null distribution reported (Supplementary Figures S4, S5, S6). New avenues are opened here for further efforts in the field of subject-level analysis.

Regarding methodological considerations, first, the optimal number of components to be derived was still a challenging question for all SS methods. The influence of this factor on the results obtained was tested using simulated data. According to real MEG data, we referred to previous experiments(O’Neill et al., 2017; Zhu et al., 2020) to set this input to 10 extracted components for all SS methods. However, not all these components are necessarily essential, especially in the case of simple tasks as motor tasks. To this end, we followed the approach of the null distribution based on sign flipping algorithm(Hunt et al., 2012; O’Neill et al., 2017; Winkler et al., 2014; Zhu et al., 2020) to select only components whose temporal dynamics significantly modulate with the task. In this way, we ensure that brain dynamics relative to the studied behavioural tasks can be summarized and described by the retained components through an automatic way that allows us to objectively compare the SS methods performance. In addition, the fact that this technique is a purely data-driven procedure that does not require any prior hypothesis or conditions manipulation makes it likely adapted to the specific examined dataset. Two points regarding significant components selection are important to mention here. First, in the applied null distribution, networks were defined to be significant if they fell outside the null distribution in either positive or negative sides because they reflect trial-onset-locked that either increases or decreases in connectivity across subjects (as amplitude envelope correlation was adopted). Second, it should be noted that components significance was evaluated relative to the specific task duration. For example, temporal duration of the entire analysis for self-paced and working memory tasks was set to be large and thus, can include dynamics that should be excluded from the analysis. Whereas, HCP motor task duration was relatively smaller so that all components extracted significantly at any time should be considered. In this context, we limited our significance interpretation/assessment in the interval of [-3;+3sec] relative to the button press instant in the case of self-paced motor task, and [-3;+16sec] relative to the visual stimulus presentation for memory task. In addition, few limitations are to be discussed when dealing with this selection. First, it was not convenient to rely on this technique when the number of trials and subjects was either too small or too big. A small number will not allow to build a reliable null distribution while a huge one will have its computational cost regarding all possible subjects’ combinations for sign-flipping procedure, as already executed in the HCP analysis. Moreover, there is no consensus about the thresholds/margins that define well a limit level for component’s amplitude. For instance, there exists networks whose temporal variation peaks at the limit of null distribution envelope. These are considered to be critical components that may be integrated in the task but considered not to be following the automatic criteria of this null distribution. Future works should therefore investigate more about this methodological approach in the framework of cognitive tasks, in addition to resting state experiments. Also, it is crucial to seek more methods to use or combine with the applied technique in order to have more robust basis for significant components selection. It is noteworthy to report that null distribution-based technique was applied uniformly for all SS methods, thus our main objective of comparison was built on a unified evaluation framework.

Second, we used the same pipeline supported by the previous studies dealing with the same dataset (cortical parcellation, source reconstruction, functional connectivity metric and source leakage correction, frequency bands and sliding window settings) (Kabbara et al., 2019; O’Neill et al., 2017). By applying already tested and validated methodological approaches, we avoid influencing factors on the comparison performed. However, we point out that other methodological solutions could be exploited by other researches using the same pipeline adopted in this work. Regarding cortical parcellation, we chose AAL atlas based on its successful use in previous MEG investigations(O’Neill et al., 2017; Tewarie et al., 2016). This atlas also provides good basis for the orthogonalisation procedure adopted since its number of regions is sufficiently low (78 ROIs) and well separated(Colclough et al., 2015). The beamformer spatial filtering was selected as the inverse problem solution due to its demonstrated efficiency in the measurement of static(Brookes et al., 2011a) and dynamic(Baker et al., 2014) functional connectivity. Functional connectivity between ROIs regions was estimated through Amplitude Envelope Correlation (AEC). This technique has been successful in elucidating electrophysiological networks of functional connectivity(Colclough et al., 2016). Other methods, such as phase couplings can be considered as an alternative way to probe different type of functional connectivity(Lachaux et al., 1999). Sliding window settings (length and step) were selected carefully as a trade-off between temporal resolution and the accuracy of the derived adjacency matrices (length=6sec, step=0.5sec for self-paced and working memory tasks)(O’Neill et al., 2017). Similarly, the same conceptual strategy was adopted for window specifications in HCP motor task to have comparable number of adjacency matrix (length=0.6sec, step=0.05sec). Concerning frequency bands, it was crucial to preprocess each dataset in its appropriate bandwidth. For example, brain signals in self-paced motor tasks were proved to be more active in the beta band, while broader range of frequency bands are integrated in complex cognitive tasks as working memory(O’Neill et al., 2017). We filtered HCP hand movements data in both beta [13-30Hz] and gamma bands [30-100Hz] separately. However, results presented in this paper refer to gamma band filtering as we noticed that motor regions were integrated much more strongly when using gamma band. This can be explained by the very rapid timescale of these executed movements which make data preprocessing in high frequencies more adequate. In this context, recent works demonstrated prominent amplitude increases in high gamma band during motor execution in motor brain regions, while beta power was enhanced after execution (delay period) (Combrisson et al., 2017; Hosaka et al., 2016). Another reason could be related to the fact that working with broader frequency range [30-100Hz] decreases the noise of adjacency matrix(O’Neill et al., 2017). Finally, to overcome source leakage problem, symmetric orthogonalisation method was applied(Colclough et al., 2015). Nevertheless, this correction method works properly only if the number of ROIs (78 in our study) was less than the degree of freedom *n* (*n = Bandwidth × window length*). This condition cannot be satisfied in the case of HCP data that deals with very small window length due to the short period time of trials. Therefore, no correction was applied with HCP data since it requires more time points.

## References

Aggarwal, C.C., Hinneburg, A., Keim, D.A., 2001. On the Surprising Behavior of Distance Metrics in High Dimensional Space, in: Van den Bussche, J., Vianu, V. (Eds.), Database Theory — ICDT 2001, Lecture Notes in Computer Science. Springer, Berlin, Heidelberg, pp. 420–434. https://doi.org/10.1007/3-540-44503-X_27

Allen, E.A., Damaraju, E., Plis, S.M., Erhardt, E.B., Eichele, T., Calhoun, V.D., 2014. Tracking WholeBrain Connectivity Dynamics in the Resting State. Cereb Cortex 24, 663–676. https://doi.org/10.1093/cercor/bhs352

Axmacher, N., Schmitz, D.P., Wagner, T., Elger, C.E., Fell, J., 2008. Interactions between Medial Temporal Lobe, Prefrontal Cortex, and Inferior Temporal Regions during Visual Working Memory: A Combined Intracranial EEG and Functional Magnetic Resonance Imaging Study. J. Neurosci. 28, 7304–7312. https://doi.org/10.1523/JNEUROSCI.1778-08.2008

Baddeley, A., Jarrold, C., Vargha-Khadem, F., 2011. Working memory and the hippocampus. J Cogn Neurosci 23, 3855–3861. https://doi.org/10.1162/jocn_a_00066

Baker, A.P., Brookes, M.J., Rezek, I.A., Smith, S.M., Behrens, T., Probert Smith, P.J., Woolrich, M., 2014. Fast transient networks in spontaneous human brain activity. eLife 3, e01867. https://doi.org/10.7554/eLife.01867

Bassett, D.S., Sporns, O., 2017. Network neuroscience. Nature neuroscience 20, 353.

Bauer, A.-K.R., Debener, S., Nobre, A.C., 2020. Synchronisation of Neural Oscillations and Cross-modal Influences. Trends in Cognitive Sciences 24, 481–495. https://doi.org/10.1016/j.tics.2020.03.003

Beckmann, C., Mackay, C., Filippini, N., Smith, S., 2009. Group comparison of resting-state FMRI data using multi-subject ICA and dual regression. NeuroImage, Organization for Human Brain Mapping 2009 Annual Meeting 47, S148. https://doi.org/10.1016/S1053-8119(09)71511-3

Bola, M., Sabel, B.A., 2015. Dynamic reorganization of brain functional networks during cognition. Neuroimage 114, 398–413.

Brookes, M.J., Hale, J.R., Zumer, J.M., Stevenson, C.M., Francis, S.T., Barnes, G.R., Owen, J.P., Morris, P.G., Nagarajan, S.S., 2011a. Measuring functional connectivity using MEG: methodology and comparison with fcMRI. Neuroimage 56, 1082–1104. https://doi.org/10.1016/j.neuroimage.2011.02.054

Brookes, M.J., Wood, J.R., Stevenson, C.M., Zumer, J.M., White, T.P., Liddle, P.F., Morris, P.G., 2011b. Changes in brain network activity during working memory tasks: a magnetoencephalography study. Neuroimage 55, 1804–1815. https://doi.org/10.1016/j.neuroimage.2010.10.074

Brookes, M.J., Woolrich, M.W., Barnes, G.R., 2012. Measuring functional connectivity in MEG: a multivariate approach insensitive to linear source leakage. Neuroimage 63, 910–920. https://doi.org/10.1016/j.neuroimage.2012.03.048

Buckner, R.L., Krienen, F.M., Castellanos, A., Diaz, J.C., Yeo, B.T.T., 2011. The organization of the human cerebellum estimated by intrinsic functional connectivity. J Neurophysiol 106, 2322–2345. https://doi.org/10.1152/jn.00339.2011

Bullmore, E., Sporns, O., 2009. Complex brain networks: graph theoretical analysis of structural and functional systems. Nat. Rev. Neurosci. 10, 186–198. https://doi.org/10.1038/nrn2575

Caminiti, S.P., Siri, C., Guidi, L., Antonini, A., Perani, D., 2015. The Neural Correlates of Spatial and Object Working Memory in Elderly and Parkinson’s Disease Subjects. Behavioural Neurology 2015, 1–10. https://doi.org/10.1155/2015/123636

Casorso, J., Kong, X., Chi, W., Van De Ville, D., Yeo, B.T.T., Liégeois, R., 2019. Dynamic mode decomposition of resting-state and task fMRI. NeuroImage 194, 42–54. https://doi.org/10.1016/j.neuroimage.2019.03.019

Cavanna, A.E., Trimble, M.R., 2006. The precuneus: a review of its functional anatomy and behavioural correlates. Brain 129, 564–583. https://doi.org/10.1093/brain/awl004

Chai, L.R., Khambhati, A.N., Ciric, R., Moore, T.M., Gur, R.C., Gur, R.E., Satterthwaite, T.D., Bassett, D.S., 2017. Evolution of brain network dynamics in neurodevelopment. Network Neuroscience 1, 14–30. https://doi.org/10.1162/NETN_a_00001

Chai, W.J., Abd Hamid, A.I., Abdullah, J.M., 2018. Working Memory From the Psychological and Neurosciences Perspectives: A Review. Front Psychol 9. https://doi.org/10.3389/fpsyg.2018.00401

Ciric, R., Nomi, J.S., Uddin, L.Q., Satpute, A.B., 2017. Contextual connectivity: A framework for understanding the intrinsic dynamic architecture of large-scale functional brain networks. Scientific Reports 7, 1–16. https://doi.org/10.1038/s41598-017-06866-w

Colclough, G.L., Brookes, M.J., Smith, S.M., Woolrich, M.W., 2015. A symmetric multivariate leakage correction for MEG connectomes. Neuroimage 117, 439–448. https://doi.org/10.1016/j.neuroimage.2015.03.071

Colclough, G.L., Woolrich, M.W., Tewarie, P.K., Brookes, M.J., Quinn, A.J., Smith, S.M., 2016. How reliable are MEG resting-state connectivity metrics? Neuroimage 138, 284–293. https://doi.org/10.1016/j.neuroimage.2016.05.070

Combrisson, E., Perrone-Bertolotti, M., Soto, J.L., Alamian, G., Kahane, P., Lachaux, J.-P., Guillot, A., Jerbi, K., 2017. From intentions to actions: Neural oscillations encode motor processes through phase, amplitude and phase-amplitude coupling. NeuroImage 147, 473–487. https://doi.org/10.1016/j.neuroimage.2016.11.042

Corbetta, M., Miezin, F.M., Dobmeyer, S., Shulman, G.L., Petersen, S.E., 1991. Selective and divided attention during visual discriminations of shape, color, and speed: functional anatomy by positron emission tomography. J. Neurosci. 11, 2383–2402.

Demb, J.B., Desmond, J.E., Wagner, A.D., Vaidya, C.J., Glover, G.H., Gabrieli, J.D., 1995. Semantic encoding and retrieval in the left inferior prefrontal cortex: a functional MRI study of task difficulty and process specificity. J. Neurosci. 15, 5870–5878. https://doi.org/10.1523/JNEUROSCI.15-09-05870.1995

Derrfuss, J., Brass, M., Yves von Cramon, D., 2004. Cognitive control in the posterior frontolateral cortex: evidence from common activations in task coordination, interference control, and working memory. Neuroimage 23, 604–612. https://doi.org/10.1016/j.neuroimage.2004.06.007

Deschamps, I., Baum, S.R., Gracco, V.L., 2014. On the role of the supramarginal gyrus in phonological processing and verbal working memory: evidence from rTMS studies. Neuropsychologia 53, 39–46. https://doi.org/10.1016/j.neuropsychologia.2013.10.015

Ding, X., Lee, J.-H., Lee, S.-W., 2013. Performance evaluation of nonnegative matrix factorization algorithms to estimate task-related neuronal activities from fMRI data. Magn Reson Imaging 31, 466–476. https://doi.org/10.1016/j.mri.2012.10.003

Dolnicar, S., n.d. A Review of Unquestioned Standards in Using Cluster Analysis for Data-Driven Market Segmentation 9.

Du, Y., Fan, Y., 2013. Group information guided ICA for fMRI data analysis. Neuroimage 69, 157–197. https://doi.org/10.1016/j.neuroimage.2012.11.008

Du, Y., Pearlson, G.D., Yu, Q., He, H., Lin, D., Sui, J., Wu, L., Calhoun, V.D., 2016. Interaction among subsystems within default mode network diminished in schizophrenia patients: A dynamic connectivity approach. Schizophr. Res. 170, 55–65. https://doi.org/10.1016/j.schres.2015.11.021

Fong, A.H.C., Yoo, K., Rosenberg, M.D., Zhang, S., Li, C.-S.R., Scheinost, D., Constable, R.T., Chun, M.M., 2019. Dynamic functional connectivity during task performance and rest predicts individual differences in attention across studies. NeuroImage 188, 14–25. https://doi.org/10.1016/j.neuroimage.2018.11.057

Fontolan, L., Morillon, B., Liegeois-Chauvel, C., Giraud, A.-L., 2014. The contribution of frequency-specific activity to hierarchical information processing in the human auditory cortex. Nat Commun 5, 4694. https://doi.org/10.1038/ncomms5694

Frackowiak, R., 1992. The anatomy of phonological and semantic processing in normal subjects. Brain, 115: 1753–1768. version 1 – 2 Dec 2008. Brain 671-682.

Friston, K.J., 1994. Functional and effective connectivity in neuroimaging: A synthesis. Human Brain Mapping 2, 56–78. https://doi.org/10.1002/hbm.460020107

Golub, G.H., Reinsch, C., n.d. Singular value decomposition and least squares solutions 18.

Grill-Spector, K., Kourtzi, Z., Kanwisher, N., 2001. The lateral occipital complex and its role in object recognition. Vision Research 41, 1409–1422. https://doi.org/10.1016/S0042-6989(01)00073-6

Grill-Spector, K., Kushnir, T., Edelman, S., Itzchak, Y., Malach, R., 1998. Cue-Invariant Activation in Object-Related Areas of the Human Occipital Lobe. Neuron 21, 191–202. https://doi.org/10.1016/S0896-6273(00)80526-7

Hassan, M., Benquet, P., Biraben, A., Berrou, C., Dufor, O., Wendling, F., 2015. Dynamic reorganization of functional brain networks during picture naming. Cortex 73, 276–288. https://doi.org/10.1016/j.cortex.2015.08.019

Hassan, M., Dufor, O., Merlet, I., Berrou, C., Wendling, F., 2014. EEG source connectivity analysis: from dense array recordings to brain networks. PloS one 9, e105041.

Hassan, M., Wendling, F., 2018. Electroencephalography Source Connectivity: Aiming for High Resolution of Brain Networks in Time and Space. IEEE Signal Process. Mag. 35, 81–96. https://doi.org/10.1109/MSP.2017.2777518

Hipp, J.F., Hawellek, D.J., Corbetta, M., Siegel, M., Engel, A.K., 2012. Large-scale cortical correlation structure of spontaneous oscillatory activity. Nat. Neurosci. 15, 884–890. https://doi.org/10.1038/nn.3101

Hosaka, R., Nakajima, T., Aihara, K., Yamaguchi, Y., Mushiake, H., 2016. The Suppression of Beta Oscillations in the Primate Supplementary Motor Complex Reflects a Volatile State During the Updating of Action Sequences. Cereb. Cortex 26, 3442–3452. https://doi.org/10.1093/cercor/bhv163

Hunt, L.T., Kolling, N., Soltani, A., Woolrich, M.W., Rushworth, M.F.S., Behrens, T.E.J., 2012. Mechanisms underlying cortical activity during value-guided choice. Nature Neuroscience 15, 470–476. https://doi.org/10.1038/nn.3017

Iraji, A., Faghiri, A., Lewis, N., Fu, Z., Rachakonda, S., Calhoun, V., 2020. Tools of the trade: Estimating time-varying connectivity patterns from fMRI data (preprint). PsyArXiv. https://doi.org/10.31234/osf.io/mvqj4

Kabbara, A., Khalil, M., O’Neill, G., Dujardin, K., El Traboulsi, Y., Wendling, F., Hassan, M., 2019. Detecting modular brain states in rest and task. Network Neuroscience 1–24.

Kabbara, A., Paban, V., Hassan, M., 2020. The dynamic modular fingerprints of the human brain at rest. bioRxiv.

Kachenoura, A., Albera, L., Senhadji, L., Comon, P., 2008. Ica: a potential tool for bci systems. IEEE Signal Process. Mag. 25, 57–68. https://doi.org/10.1109/MSP.2008.4408442

Kourtzi, Z., Kanwisher, N., 2001. Representation of perceived object shape by the human lateral occipital complex. Science 293, 1506–1509. https://doi.org/10.1126/science.1061133

Lachaux, J.-P., Rodriguez, E., Martinerie, J., Varela, F.J., 1999. Measuring phase synchrony in brain signals. Human Brain Mapping 8, 194–208. https://doi.org/10.1002/(SICI)1097-0193(1999)8:4<194::AID-HBM4>3.0.CO;2-C

Lee, D.D., Seung, H.S., 1999. Learning the parts of objects by non-negative matrix factorization. Nature 401, 788–791. https://doi.org/10.1038/44565

Leonardi, N., n.d. Dynamic brain networks explored by structure-revealing methods 170.

Liu, X., Duyn, J.H., 2013. Time-varying functional network information extracted from brief instances of spontaneous brain activity. PNAS 110, 4392–4397. https://doi.org/10.1073/pnas.1216856110

Lloyd, S., 1982. Least squares quantization in PCM. IEEE Trans. Inform. Theory 28, 129–137. https://doi.org/10.1109/TIT.1982.1056489

Melnik, A., Hairston, W.D., Ferris, D.P., König, P., 2017. EEG correlates of sensorimotor processing: independent components involved in sensory and motor processing. Scientific Reports 7, 4461. https://doi.org/10.1038/s41598-017-04757-8

Metzak, P., Feredoes, E., Takane, Y., Wang, L., Weinstein, S., Cairo, T., Ngan, E.T.C., Woodward, T.S., 2011. Constrained principal component analysis reveals functionally connected load-dependent networks involved in multiple stages of working memory. Hum Brain Mapp 32, 856–871. https://doi.org/10.1002/hbm.21072

Meyer-Baese, A., Wismueller, A., Lange, O., 2004. Comparison of two exploratory data analysis methods for fMRI: unsupervised clustering versus independent component analysis. IEEE Trans Inf Technol Biomed 8, 387–398. https://doi.org/10.1109/titb.2004.834406

Mheich, A., Hassan, M., Khalil, M., Berrou, C., Wendling, F., 2015. A new algorithm for spatiotemporal analysis of brain functional connectivity. Journal of Neuroscience Methods 242, 77–81. https://doi.org/10.1016/j.jneumeth.2015.01.002

Mheich, A., Hassan, M., Khalil, M., Gripon, V., Dufor, O., Wendling, F., 2018. SimiNet: A Novel Method for Quantifying Brain Network Similarity. IEEE Transactions on Pattern Analysis and Machine Intelligence 40, 2238–2249. https://doi.org/10.1109/TPAMI.2017.2750160

Miller, R.L., Yaesoubi, M., Turner, J.A., Mathalon, D., Preda, A., Pearlson, G., Adali, T., Calhoun, V.D., 2016. Higher Dimensional Meta-State Analysis Reveals Reduced Resting fMRI Connectivity Dynamism in Schizophrenia Patients. PLOS ONE 11, e0149849. https://doi.org/10.1371/journal.pone.0149849

Mion, M., Patterson, K., Acosta-Cabronero, J., Pengas, G., Izquierdo-Garcia, D., Hong, Y.T., Fryer, T.D., Williams, G.B., Hodges, J.R., Nestor, P.J., 2010. What the left and right anterior fusiform gyri tell us about semantic memory. Brain 133, 3256–3268. https://doi.org/10.1093/brain/awq272

Negrón-Oyarzo, I., Espinosa, N., Aguilar-Rivera, M., Fuenzalida, M., Aboitiz, F., Fuentealba, P., 2018. Coordinated prefrontal-hippocampal activity and navigation strategy-related prefrontal firing during spatial memory formation. Proc. Natl. Acad. Sci. U.S.A. 115, 7123–7128. https://doi.org/10.1073/pnas.1720117115

Nestor, P.J., Fryer, T.D., Hodges, J.R., 2006. Declarative memory impairments in Alzheimer’s disease and semantic dementia. Neuroimage 30, 1010–1020. https://doi.org/10.1016/j.neuroimage.2005.10.008

O’Neill, G.C., Bauer, M., Woolrich, M.W., Morris, P.G., Barnes, G.R., Brookes, M.J., 2015. Dynamic recruitment of resting state sub-networks. NeuroImage 115, 85–95. https://doi.org/10.1016/j.neuroimage.2015.04.030

O’Neill, G.C., Tewarie, P.K., Colclough, G.L., Gascoyne, L.E., Hunt, B.A.E., Morris, P.G., Woolrich, M.W., Brookes, M.J., 2017. Measurement of dynamic task related functional networks using MEG. NeuroImage 146, 667–678. https://doi.org/10.1016/j.neuroimage.2016.08.061

Pomper, U., Keil, J., Foxe, J.J., Senkowski, D., 2015. Intersensory selective attention and temporal orienting operate in parallel and are instantiated in spatially distinct sensory and motor cortices: Human Intersensory and Temporal Attention. Hum. Brain Mapp. 36, 3246–3259. https://doi.org/10.1002/hbm.22845

Rouhinen, S., Siebenhühner, F., Palva, J.M., Palva, S., 2020. Spectral and Anatomical Patterns of Large-Scale Synchronization Predict Human Attentional Capacity. Cereb. Cortex. https://doi.org/10.1093/cercor/bhaa110

Sahonero-Alvarez, G., Calderon, H., 2017. A Comparison of SOBI, FastICA, JADE and Infomax Algorithms 6.

Tewarie, P., Bright, M.G., Hillebrand, A., Robson, S.E., Gascoyne, L.E., Morris, P.G., Meier, J., Van Mieghem, P., Brookes, M.J., 2016. Predicting haemodynamic networks using electrophysiology: The role of non-linear and cross-frequency interactions. Neuroimage 130, 273–292. https://doi.org/10.1016/j.neuroimage.2016.01.053

Tewarie, P., Hunt, B.A.E., O’Neill, G.C., Byrne, A., Aquino, K., Bauer, M., Mullinger, K.J., Coombes, S., Brookes, M.J., 2019. Relationships Between Neuronal Oscillatory Amplitude and Dynamic Functional Connectivity. Cereb. Cortex 29, 2668–2681. https://doi.org/10.1093/cercor/bhy136

Thomas Yeo, B.T., Krienen, F.M., Sepulcre, J., Sabuncu, M.R., Lashkari, D., Hollinshead, M., Roffman, J.L., Smoller, J.W., Zöllei, L., Polimeni, J.R., Fischl, B., Liu, H., Buckner, R.L., 2011. The organization of the human cerebral cortex estimated by intrinsic functional connectivity. Journal of Neurophysiology 106, 1125–1165. https://doi.org/10.1152/jn.00338.2011

van den Heuvel, M.P., Hulshoff Pol, H.E., 2010. Exploring the brain network: A review on resting-state fMRI functional connectivity. European Neuropsychopharmacology 20, 519–534. https://doi.org/10.1016/j.euroneuro.2010.03.008

Van Essen, D.C., Ugurbil, K., Auerbach, E., Barch, D., Behrens, T.E.J., Bucholz, R., Chang, A., Chen, L., Corbetta, M., Curtiss, S.W., Della Penna, S., Feinberg, D., Glasser, M.F., Harel, N., Heath, A.C., Larson-Prior, L., Marcus, D., Michalareas, G., Moeller, S., Oostenveld, R., Petersen, S.E., Prior, F., Schlaggar, B.L., Smith, S.M., Snyder, A.Z., Xu, J., Yacoub, E., WU-Minn HCP Consortium, 2012. The Human Connectome Project: a data acquisition perspective. Neuroimage 62, 2222–2231. https://doi.org/10.1016/j.neuroimage.2012.02.018

Vandenberghe, R., Price, C., Wise, R., Josephs, O., Frackowiak, R.S.J., 1996. Functional anatomy of a common semantic system for words and pictures. Nature 383, 254–256. https://doi.org/10.1038/383254a0

Vidaurre, D., Abeysuriya, R., Becker, R., Quinn, A.J., Alfaro-Almagro, F., Smith, S.M., Woolrich, M.W., 2018a. Discovering dynamic brain networks from big data in rest and task. NeuroImage, Brain Connectivity Dynamics 180, 646–656. https://doi.org/10.1016/j.neuroimage.2017.06.077

Vidaurre, D., Hunt, L.T., Quinn, A.J., Hunt, B.A.E., Brookes, M.J., Nobre, A.C., Woolrich, M.W., 2018b. Spontaneous cortical activity transiently organises into frequency specific phase-coupling networks. Nature Communications 9, 2987. https://doi.org/10.1038/s41467-018-05316-z

Vidaurre, D., Quinn, A.J., Baker, A.P., Dupret, D., Tejero-Cantero, A., Woolrich, M.W., 2016. Spectrally resolved fast transient brain states in electrophysiological data. NeuroImage 126, 81–95. https://doi.org/10.1016/j.neuroimage.2015.11.047

Vigneau, M., Beaucousin, V., Hervé, P.Y., Duffau, H., Crivello, F., Houdé, O., Mazoyer, B., Tzourio-Mazoyer, N., 2006. Meta-analyzing left hemisphere language areas: phonology, semantics, and sentence processing. Neuroimage 30, 1414–1432. https://doi.org/10.1016/j.neuroimage.2005.11.002

Wilkins, K.B., Yao, J., 2020. Coordination of multiple joints increases bilateral connectivity with ipsilateral sensorimotor cortices. Neuroimage 207, 116344. https://doi.org/10.1016/j.neuroimage.2019.116344

Winkler, A.M., Ridgway, G.R., Webster, M.A., Smith, S.M., Nichols, T.E., 2014. Permutation inference for the general linear model. Neuroimage 92, 381–397. https://doi.org/10.1016/j.neuroimage.2014.01.060

Xie, J., Douglas, P.K., Wu, Y.N., Brody, A.L., Anderson, A.E., 2017. Decoding the encoding of functional brain networks: An fMRI classification comparison of non-negative matrix factorization (NMF), independent component analysis (ICA), and sparse coding algorithms. J. Neurosci. Methods 282, 81–94. https://doi.org/10.1016/j.jneumeth.2017.03.008

Yaesoubi, M., Miller, R.L., Calhoun, V.D., 2015. Mutually temporally independent connectivity patterns: A new framework to study the dynamics of brain connectivity at rest with application to explain group difference based on gender. NeuroImage 107, 85–94. https://doi.org/10.1016/j.neuroimage.2014.11.054

Yamashita, M., Kawato, M., Imamizu, H., 2015. Predicting learning plateau of working memory from whole-brain intrinsic network connectivity patterns. Sci Rep 5, 1–8. https://doi.org/10.1038/srep07622

Yousry, T., 1997. Localization of the motor hand area to a knob on the precentral gyrus. A new landmark. Brain 120, 141–157. https://doi.org/10.1093/brain/120.1.141

Zhu, Y., Liu, J., Ye, C., Mathiak, K., Astikainen, P., Ristaniemi, T., Cong, F., 2020. Discovering dynamic task-modulated functional networks with specific spectral modes using MEG. NeuroImage 218, 116924. https://doi.org/10.1016/j.neuroimage.2020.116924

